# On the origin and evolution of RNA editing in metazoans

**DOI:** 10.1101/2020.01.19.911685

**Authors:** Qiye Li, Pei Zhang, Ji Li, Hao Yu, Xiaoyu Zhan, Yuanzhen Zhu, Qunfei Guo, Huishuang Tan, Nina Lundholm, Lydia Garcia, Michael D. Martin, Meritxell Antó Subirats, Yi-Hsien Su, Iñaki Ruiz-Trillo, Mark Q. Martindale, Jr-Kai Yu, M. Thomas P. Gilbert, Guojie Zhang

**Affiliations:** BGI-Shenzhen, Shenzhen 518083, China; State Key Laboratory of Genetic Resources and Evolution, Kunming Institute of Zoology, Chinese Academy of Sciences, Kunming 650223, China; Section for Ecology and Evolution, Department of Biology, University of Copenhagen, DK-2100 Copenhagen, Denmark; China National Genebank, BGI-Shenzhen, Shenzhen 518120, China; School of Basic Medicine, Qingdao University, Qingdao 266071, China; College of Life Science and Technology, Huazhong University of Science and Technology, Wuhan 430074, China; Center for Informational Biology, University of Electronic Science and Technology of China, Chengdu 611731, China; Natural History Museum of Denmark, University of Copenhagen, Copenhagen 1350, Denmark; Department of Natural History, NTNU University Museum, Norwegian University of Science and Technology (NTNU), NO-7491 Trondheim, Norway; Center for Theoretical Evolutionary Genomics, Dept. of Integrative Biology, University of California Berkeley, Berkeley, California 94720, USA; Institute of Evolutionary Biology, UPF-CSIC Barcelona, 08003 Barcelona, Spain; Institute of Cellular and Organismic Biology, Academia Sinica, 11529 Taipei, Taiwan; ICREA, Passeig Lluís Companys 23, 08010 Barcelona, Catalonia, Spain; Departament de Genètica, Microbiologia i Estadística, Facultat de Bilogia, Universitat de Barcelona (UB), Barcelona 08028, Spain; The Whitney Laboratory for Marine Bioscience, University of Florida, St. Augustine, FL 32080, USA; Marine Research Station, Institute of Cellular and Organismic Biology, Academia Sinica, 26242 Yilan, Taiwan; Section for Evolutionary Genomics, The GLOBE Institute, University of Copenhagen, Copenhagen 1352, Denmark; Center for Excellence in Animal Evolution and Genetics, Chinese Academy of Sciences, 650223, Kunming, China

## Abstract

Extensive adenosine-to-inosine (A-to-I) editing of nuclear-transcribed RNAs is the hallmark of metazoan transcriptional regulation, and is fundamental to numerous biochemical processes. Here we explore the origin and evolution of this regulatory innovation, by quantifying its prevalence in 22 species that represent all major transitions in metazoan evolution. We provide substantial evidence that extensive RNA editing emerged in the common ancestor of extant metazoans. We find the frequency of RNA editing varies across taxa in a manner independent of metazoan complexity. Nevertheless, cis-acting features that guide A-to-I editing are under strong constraint across all metazoans. RNA editing seems to preserve an ancient mechanism for suppressing the more recently evolved repetitive elements, and is generally nonadaptive in protein-coding regions across metazoans, except for *Drosophila* and cephalopods. Interestingly, RNA editing preferentially target genes involved in neurotransmission, cellular communication and cytoskeleton, and recodes identical amino acid positions in several conserved genes across diverse taxa, emphasizing broad roles of RNA editing in cellular functions during metazoan evolution that have been previously underappreciated.

## Introduction

The central dogma of molecular biology emphasizes how genetic information passes faithfully from DNA, to RNA, to proteins. However, this dogma has been challenged by the phenomenon of RNA editing — a post/co-transcriptional-processing mechanism that can alter RNA sequences by insertion, deletion or substitution of specific nucleotides, thus producing transcripts that are not directly encoded in the genome ^1^. In metazoans, the most prevalent form of RNA editing is the deamination of adenosine (A) to inosine (I), which is catalyzed by a family of adenosine deaminases acting on RNA (ADARs) ^2, 3^. As inosine is recognized *in vivo* as guanosine by ribosomes and other molecular machinery, RNA editing can affect almost all aspects of cellular RNA functions, from changing mRNA coding potential by altering codons or splicing patterns, to regulating the cellular fate of mRNA by editing its microRNA (miRNA) binding sites ^4–6^. RNA editing is particularly pervasive in neural systems, where it has been shown to modulate neural development processes ^7, 8^, neural network plasticity ^9, 10^ and organismal adaptation to environmental changes ^11–13^. Defects in RNA editing machinery have been linked to a variety of neurological diseases, autoimmune disorders and cancers ^14–18^.

Although recent high-throughput sequencing-based analyses have identified a surprisingly large number of RNA-editing sites in different metazoans, including humans ^19–23^, mice ^24, 25^, *Caenorhabditis elegans* ^26^, fruit flies ^27–30^, ants ^31^, bumblebees ^32^ and cephalopods ^33, 34^, conclusions about the evolutionary patterns of this phenomenon are inconsistent. For example, while almost all human RNA-editing sites occur in Alu repeat elements ^20, 21^, editing in *Drosophila* primarily targets exonic (particularly coding) regions ^27, 28^. Additionally, while recoding RNA editing, which leads to nonsynonymous substitutions in protein-coding sequences, is abundant and affects around half of the protein-coding genes in coleoid cephalopods ^33, 34^, it is relatively rare in mammals and insects ^21, 24, 28, 31, 32^. Furthermore, while recoding editing in humans is generally nonadaptive ^35^, it is typically adaptive in *Drosophila* and cephalopods ^28, 34^. More importantly, although the *ADAR* gene family is considered to have originated in the common ancestor of extant metazoans ^36^, the functional activity of ADARs in catalyzing RNA editing in most metazoan lineages actually remains unknown, especially in those earliest branching lineages like Ctenophora and Porifera.

In summary, many fundamental questions about the nature of metazoan RNA editing remain to be investigated, including: When did RNA editing emerge during metazoan evolution? Are there conserved sequence features that underly RNA editing in all metazoans? What genes and genomic elements are the primary targets of metazoan RNA editing? How does the prevalence of recoding editing vary by lineage, and does it generally provide adaptive amino acid changes in metazoans? Addressing these questions requires the characterization of RNA editomes across the diversity of metazoans and their closest unicellular relatives, thus we systematically investigated the prevalence and characteristics of RNA editing in 22 lineages that encompass the key transitions in metazoan evolution.

## Results

### Profiling the RNA editomes across the phylogeny of metazoans

We performed both DNA-seq and strand-specific RNA-seq for 18 species, including 14 metazoans and 4 unicellular eukaryotes closely related to animals. 14 out of these 18 species have not been subjected to transcriptome-wide RNA editing investigation previously (Fig. 1a). For each species, two to three (mostly three) biological replicates were sequenced, yielding 3.27 Tbp (tera base pairs) sequencing data in total, with the average DNA and RNA coverage achieving 75X (ranging 15-345X) and 45X (ranging 10-162X) respectively for each biological replicate after alignment (Supplementary Table 1). Together with published sequencing data from *C. elegans* ^26^, ant ^31^, octopus ^37^ and human ^22^ (Supplementary Table 1), we were able to profile and compare the RNA editomes of 22 species, which represent nearly all the major phyla of extant metazoans, including the earliest-branching lineages Ctenophora, Porifera and Placozoa, as well as their closest unicellular relatives Choanoflagellatea, Filasterea and Ichthyosporea (Fig. 1a). These data thus provide the first opportunity to phylogenetically investigate the prevalence of RNA editing within Holozoa, the clade that includes animals and their closest single-celled relatives ^38^.

**Figure 1.**
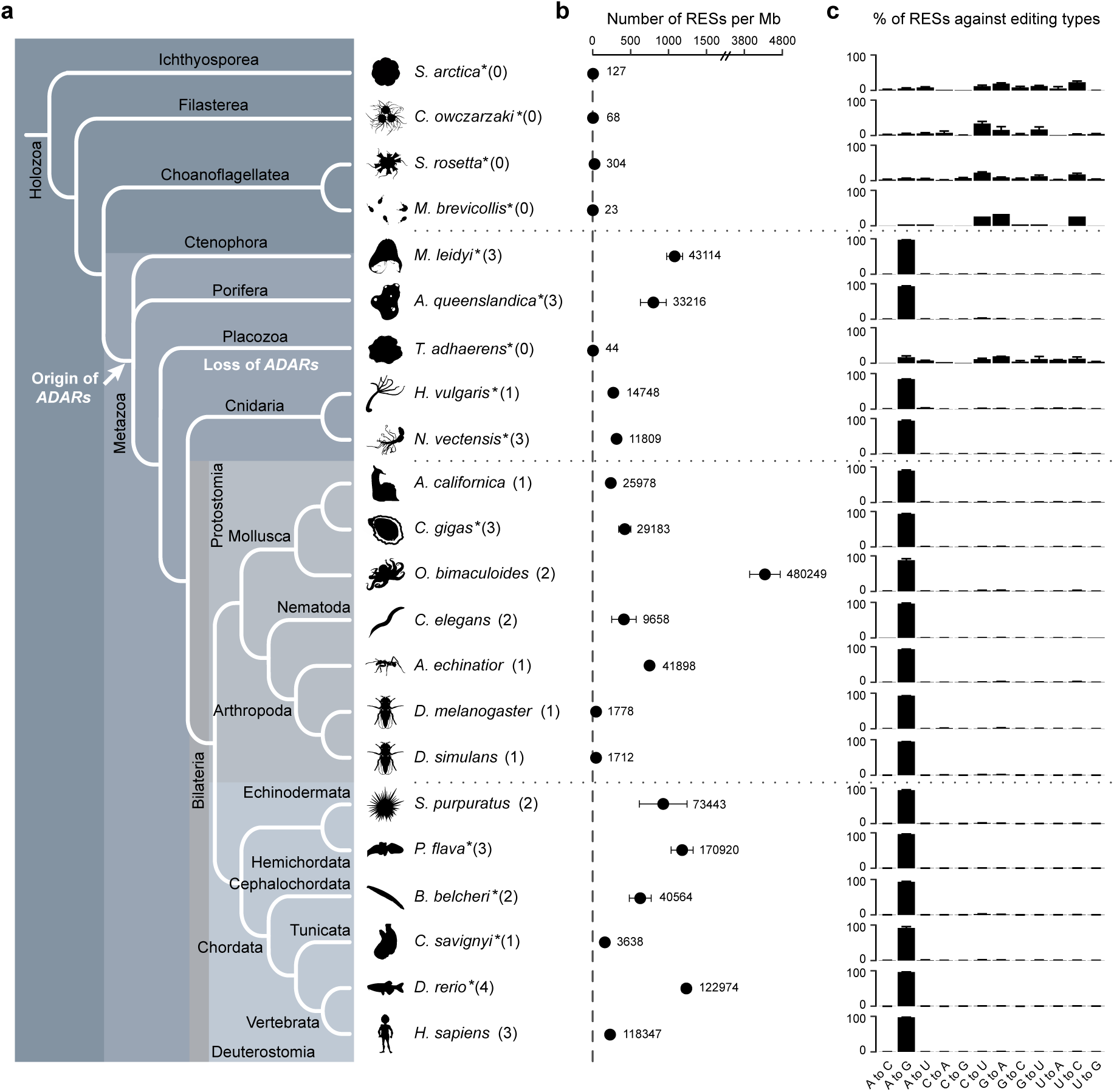
The origin and variation of RNA editing in metazoans. (**a**) The phylogeny of the 22 species examined in this study, with the inferred origin and lineage-specific loss of *ADARs* indicated. The topology of the phylogenetic tree is derived according to previous reports ^87–89^. The copy number of *ADAR-like* genes identified in the genome of each species is present in parenthesis after the Latin name. Asterisks (*) indicate species that have not previously been subject to transcriptome-wide RNA editing analyses. Full names for the 22 species from top to bottom are *Sphaeroforma arctica* (ichthyosporean), *Capsaspora owczarzaki* (filasterean), *Salpingoeca rosetta* (choanoflagellate), *Monosiga brevicollis* (choanoflagellate), *Mnemiopsis leidyi* (comb jelly), *Amphimedon queenslandica* (sponge), *Trichoplax adhaerens* (placozoan), *Hydra vulgaris* (hydra), *Nematostella vectensis* (sea anemone), *Aplysia californica* (sea hare), *Crassostrea gigas* (oyster), *Octopus bimaculoides* (octopus), *Caenorhabditis elegans* (roundworm), *Acromyrmex echinatior* (ant), *Drosophila melanogaster* (fruit fly), *Drosophila simulans* (fruit fly), *Strongylocentrotus purpuratus* (sea urchin), *Ptychodera flava* (acorn worm), *Branchiostoma belcheri* (lancelet), *Ciona savignyi* (sea squirt), *Danio rerio* (zebrafish) and *Homo sapiens* (human). (**b**) The occurrence rate of RNA editing in each species, which was measured as the number of RNA-editing sites per million transcribed genomic sites (RNA depth ≥ 2X). The mean number of editing sites identified in each species is presented on the right of each dot. (**c**) Percentage of editing sites across the 12 possible types of nucleotide substitutions. Error bars in panels **b** and **c** represent s.d. across samples.

Given that some RNA-editing sites tend to appear in clusters, while others remain isolated, we adopted two complementary methods to identify the RNA editomes for each species. Briefly, we first employed RES-Scanner ^39^ to identify RNA-editing sites by comparing the matching DNA- and RNA-seq data from the same sample. This method has high accuracy when searching for RNA-editing sites that are isolated or not heavily clustered. We next performed hyper-editing detection ^40^, using the RNA reads that failed to align by RES-Scanner, in order to capture the hyper-edited reads and the clusters of editing sites they harbored. The results of RES-Scanner and hyper-editing detection were combined to yield the RNA editome of each sample (Supplementary Table 2). We have compiled the whole pipeline as an easy-to-use software package named RES-Scanner2, which is applicable to transcriptome-wide identification of RNA-editing sites in any species with matching DNA- and RNA-seq data (see Methods for details).

### Extensive RNA editing emerged in the last common ancestor of modern metazoans accompanied by the origin of *ADARs*

We detected very few putative RNA-editing sites (ranging 23-304) in the four unicellular holozoans (Fig. 1b and Supplementary Table 2). No dominant type of nucleotide substitution was observed (Fig. 1c), and the frequency of each type of nucleotide substitution was close to that of genetic polymorphism (Supplementary Fig. 1a), implying that RNA-editing sites detected in these species likely represent noise. In contrast thousands, to hundreds of thousands, of RNA-editing sites were identified in almost all the sampled metazoans, including the earliest-branching Ctenophora and Porifera, with the vast majority (>90%) consisting of A-to-G substitutions (i.e. A-to-I editing; Fig. 1b,c). The only exception was *Trichoplax adhaerens*, a morphology-simplified metazoan belonging to Placozoa (a sister group to Cnidaria and Bilateria) ^41^. Concordantly, we confirmed the existence of *ADAR-like* genes in all the sampled species except *T. adhaerens* and the unicellular taxa (Fig. 1a and Supplementary Table 3; See Methods). Our results thus provide direct evidence that extensive editing of nuclear-transcribed RNAs first emerged in the last common ancestor of modern metazoans, alongside the appearance of ADAR-mediated A-to-I editing, which is pervasively preserved in most extant animal lineages. We also highlight that our detection methods do not depend on any prior knowledge about the dominate type of RNA editing in any species studied (see Methods), thus, our results also imply that RNA editing in any manner other than A-to-I, is either extremely rare, or non-existent, in the animal kingdom.

We next calculated the occurrence rate of RNA editing per genome by counting the number of RNA-editing sites per million transcribed genomic sites (i.e. sites with RNA depth ≥ 2X). Our results indicate that the octopus exhibits the highest, and *Drosophila* the lowest, number and occurrence rate among the sampled taxa that have the RNA-editing machinery. Surprisingly, the occurrence rates in the ctenophore *Mnemiopsis leidyi* and sponge *Amphimedon queenslandica* are higher than that of all sampled cnidarians and many bilaterians (Fig. 1b), while humans are among the species with lowest rates (Fig. 1b). Similar patterns were obtained if we weighted each editing site with its editing level, or if we only considered A-to-I editing (Supplementary Fig. 1b-e). These results suggest that the global level of RNA editing has changed considerably during the diversification of metazoan, and does not increase directly alongside organismal complexity.

### The A-to-I editing associated sequence features are under strong constraint in metazoans

Consistent with the double-stranded RNA (dsRNA) binding property of ADAR enzymes ^2, 6^, we observed that A-to-I editing sites in all the sampled metazoans with *ADARs* were preferentially located in potential dsRNA regions that could form by intramolecular folding of pre-mRNA. Specifically, we found on average that 37% (ranging 6% to 86%) of the editing sites target regions that show a reverse-complement alignment within their upstream or downstream sequences, which is significantly higher than the expected levels of ∼1% calculated from randomly selected transcribed adenosines (Fig. 2a; See methods). These results confirm that a stable dsRNA structure is critical for establishing A-to-I editing *in vivo* across metazoans ^42^, and further reveal that intramolecular folding of pre-mRNA is a major way to form dsRNA substrates for A-to-I editing in most species.

**Figure 2.**
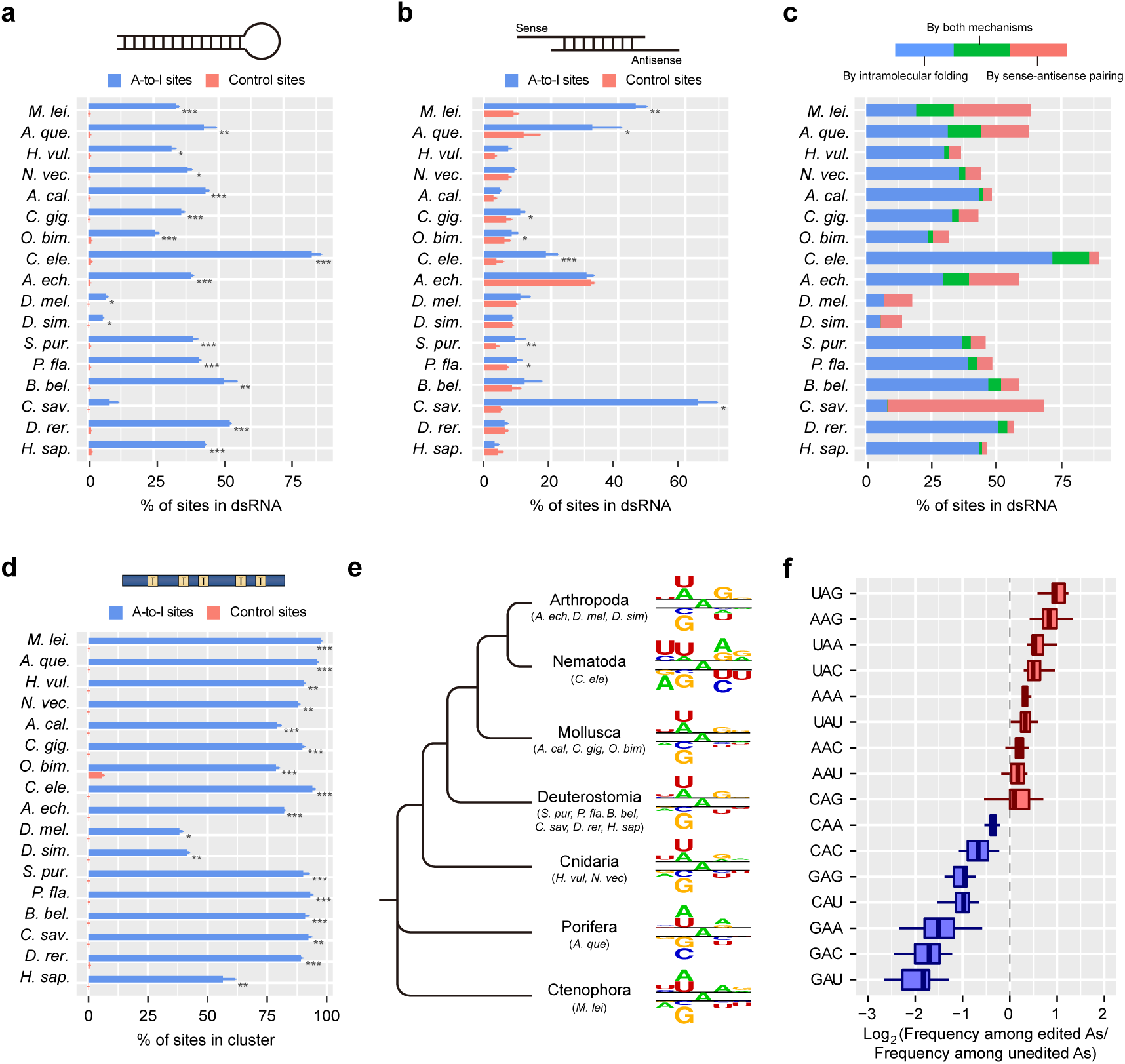
The structure and sequence preferences of A-to-I editing in metazoans. (**a**) Percentage of A-to-I editing sites locating in dsRNA regions potentially formed by intramolecular folding of pre-mRNA, measured as the proportion of sites locating in a region (± 200 nt centered on the edited adenosine) that shows a reverse-complement alignment (identity ≥ 80% and aligned length ≥ 50 nt) within its flanking sequences (± 2 knt centered on the edited adenosine). Control sites were randomly selected transcribed adenosines with the same number and comparable RNA depth of the A-to-I editing sites in each sample of each species (see Methods for details). (**b**) Percentage of A-to-I editing sites locating in dsRNA regions potentially formed by intermolecular hybridization of sense-antisense transcripts, measured as the proportion of sites locating in a region (± 50 nt centered on the edited adenosine) with transcription signal (RNA depth ≥ 2X along >50% of the region) in the both strands. Control sites were the same set of adenosine sites used in panel **a**. (**c**) Percentage of A-to-I editing sites locating in dsRNA regions, either potentially formed by intramolecular folding of pre-mRNA, or by intermolecular hybridization of sense-antisense transcripts or by both mechanisms. (**d**) Percentage of A-to-I editing sites occurring in clusters. A cluster contains ≥ 3 A-to-I editing sites of which the distance for two adjacent sites ≤ 30 nt. Control sites were the same set of adenosine sites used in panel **a**. (**e**) Neighboring nucleotide preference of the edited adenosines in comparison to the unedited transcribed adenosines within ± 50 nt of the edited adenosines. For the lineages with more than one representative species, the same number of editing sites were randomly selected according to the species with the lowest number of editing sites in this lineage. Nucleotides were plotted using the size of the nucleotide that was proportional to the difference between the edited and unedited datasets, with the upper part presenting enriched nucleotides in the edited dataset and lower part presenting depleted nucleotides. (**f**) The relative frequency of the 16 nucleotide triplets with adenosine in the center subject to A-to-I editing, measured as the frequency of a triplet observed among all edited adenosines against the frequency of this triplet observed among all neighboring unedited adenosines within ± 50 nt of the edited adenosines. Boxplot shows the distribution across the 17 metazoans with *ADARs*, and nucleotide triplets are ordered according to highest median value across species to the lowest. Error bars in panels **a, b** and **d** represent s.d. across samples, and asterisks indicate significance levels estimated by two-tailed paired t-tests with “*” representing *p* < 0.05, “**” < 0.01 and “***” < 0.001.

Intermolecular hybridization of sense and antisense transcripts is another potential mechanism to form dsRNA ^43^, but its role in inducing A-to-I editing is thought to be negligible in mammals^44^. Taking advantage of the strand information provided by strand-specific RNA-seq, we found that the proportions of editing sites that were located in regions containing transcription signals on both strands (mean 17%, ranging 3% to 64%) were significantly higher than the expected levels (mean 8%, ranging 3% to 32%) in 8 out of the 17 metazoans with *ADARs* (Fig. 2b; See methods). In particular, while for most species there are generally many more editing sites found in potential dsRNA regions formed by intramolecular folding, the ctenophore *M. leidyi* and sea squirt *Ciona savignyi* showed a reverse tendency, with higher proportions of editing sites found in regions with transcription signals in both strands (Fig. 2c). This implies that intermolecular hybridization of sense and antisense transcripts likely represents an important means for forming dsRNA substrates for A-to-I editing, in at least some taxa. This conclusion is further supported by the significantly higher-than-expected proportion of A-to-I editing sites locating in regions targeted by RNA editing on both strands in many species (Supplementary Fig. 2a,c).

With regards to the genomic distribution of A-to-I editing, we found on average 81% (ranging 41% to 97%) of the metazoan editing sites were clustered, which is significantly higher than the expected levels of less than l% (Fig. 2d). The median distances between any two adjacent editing sites were mostly around 5 nt (ranging 4 to 81 nt; Supplementary Table 4). Furthermore, editing levels of the clustered editing sites were generally higher than those of isolated sites, except in *Hydra vulgaris*, *Drosophila*, *C. savignyi* and humans (Supplementary Fig. 2b). A typical metazoan editing cluster (i.e. a region with ≥ 3 A-to-I editing sites and the distance of two adjacent sites ≤ 30 nt) was ∼50 nt in length, and harbored 9 A-to-I editing sites, and we estimated that up to 52% of the adenosines within a cluster were targeted by RNA editing (Supplementary Table 4). Taken together, our results indicate that the majority of metazoan A-to-I editing sites are organized in dense clusters, within RNA regions that can form stable dsRNA structures.

Since ADARs recognize dsRNA when exerting A-to-I editing, we then asked what primary sequence motifs guide ADARs to preferentially edit certain adenosines rather than others in their dsRNA substrates. By comparing the surrounding sequence context of edited adenosine sites to neighboring unedited adenosine sites (i.e. unedited adenosines with RNA depth ≥ 2X and within ± 50 nt of the edited adenosines), we observed clear and conserved nucleotide preferences for the positions that are directly 5’ and 3’ adjacent to the edited adenosines (i.e. the -1 and +1 positions). Specifically, the 5’ adjacent position strongly favored uridine and adenosine, but disfavored guanosine across all metazoans, and to a lesser extent, cytosine was also disfavored (Fig. 2e and Supplementary Fig. 3). In contrast, the nucleotide preference for the 3’ adjacent position is relatively weaker, and less conserved, with guanosine being favored and uridine being disfavored in most species (Fig. 2e and Supplementary Fig. 3). This implies that the 5’ adjacent position has the most influential and a conserved role on determining whether an adenosine will be edited. Concordantly, we found the nucleotide triplets of U<u>A</u>G and A<u>A</u>G, with the edited adenosines in the center, to be the most likely edited triplets, while G<u>A</u>U was the least likely edited triplet in metazoans (Fig. 2f).

Interestingly, *C. elegans* also displayed a strong sequence preference for the 5’ second nearest (-2) position of the edited adenosines that is not observed in other metazoans, with uridine being strongly favored and adenosine being strongly disfavored (Fig. 2e and Supplementary Fig. 3). We speculate that this *C. elegans* specific motif adjustment is associated with the high sequence divergence of the *C. elegans* ADARs against other metazoan ADARs, as phylogenetic analyses separate both the *C. elegans* ADR-1 and ADR-2 from other metazoans (Supplementary Fig. 2d), and both *C. elegans* ADR-1 and ADR-2 show high nonsynonymous substitution rates (*d_N_*) against ADARs from other metazoans (Supplementary Fig. 2e).

### Evolutionarily young repetitive elements are the primary targets of metazoan RNA editing

In all metazoans sampled except for the two fruit flies and sea squirt, repetitive elements including transposons and tandem repeats were the major targets of A-to-I editing, and harbored on average 83% (ranging 73% to 95%) of the editing sites (Fig. 3a). This suggests that extensive editing of repeat-containing transcripts is the ancestral and predominant feature for metazoan RNA editing, probably because these regions are more likely to hybridize with nearby oppositely oriented repeats, creating the dsRNA structures suitable for ADARs binding (Supplementary Fig. 4c). It is noteworthy that, even for those sites on pre-mRNA (i.e. exon + intron) of protein-coding genes, especially those outside coding regions, the majority (>70%) were also associated with repetitive elements (Supplementary Fig. 4d). This implies that most editable sites on protein-coding genes were actually introduced by the invasion of repetitive elements into gene regions.

**Figure 3.**
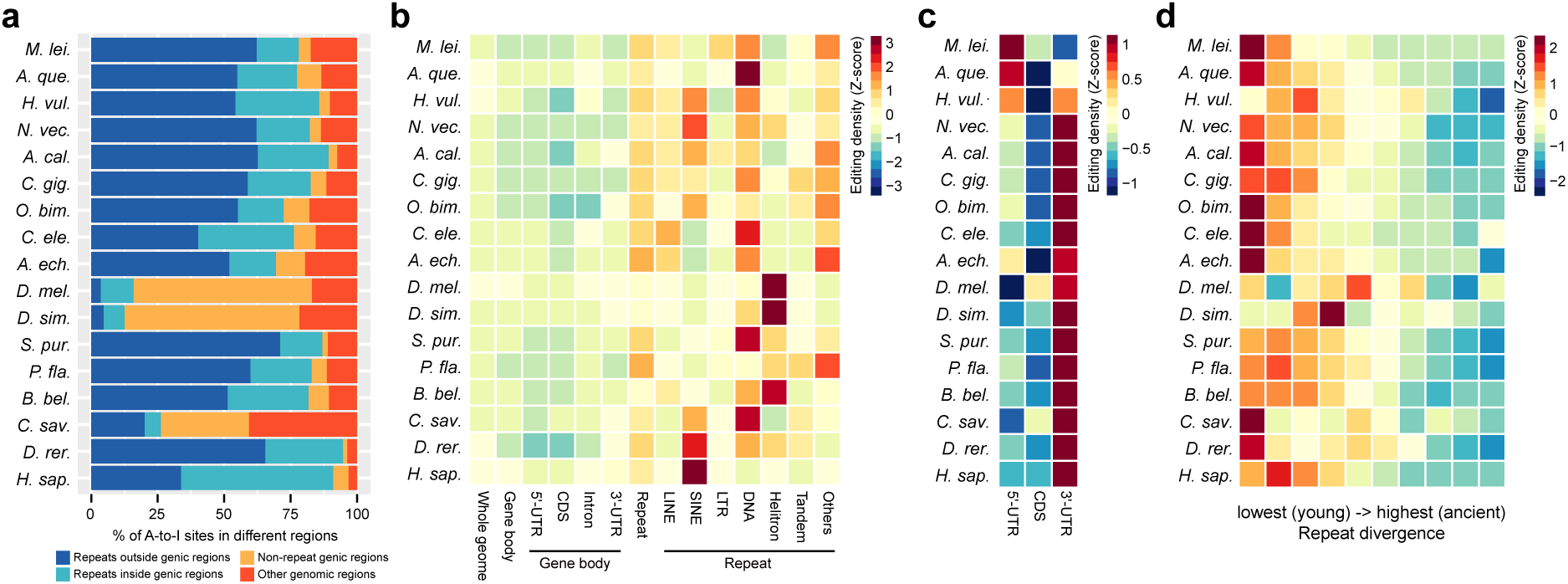
The primary genomic targets of metazoan A-to-I editing. (**a**) The proportion of A-to-I editing sites versus total A-to-I editing sites in different genomic regions. Genic regions include untranslated regions (5’-UTR and 3’UTR), CDS and intron of all protein-coding genes. Repeats include transposons and tandem repeats annotated for each species in this study (see Methods). (**b**) Comparison of A-to-I editing density across different genomic elements in each species. Editing density of an element was calculated as the number of A-to-I editing sites locating in this element divided by the number of transcribed adenosines (RNA depth ≥ 2X) in this element. (**c**) Comparison of editing density across 5’-UTR, CDS and 3’-UTR. Note the different scales used between panel **b** and **c** in order to show the difference of editing density among genic elements. (**d**) The negative correlation between the sequence divergence and editing density of repetitive elements.

Given that the total lengths of the different genomic elements vary greatly within each genome, we next calculated the A-to-I editing density for each type of genomic element, by counting the number of editing sites per million of transcribed adenosine sites (i.e. RNA depth ≥ 2X). After this normalization, we observed that the editing densities of protein-coding gene-related elements (i.e. 5’-UTR, CDS, intron and 3’-UTR) were close to the whole genome average level in all metazoans (Fig. 3b). However, editing densities generally increased from 5’ to 3’ of mRNA transcripts, with 3’ UTR being relatively more favored by A-to-I editing than 5’-UTR and CDS (Fig. 3c), consistent with previous observation in *Drosophila* ^27, 28^. In contrast, the editing densities of repetitive elements, especially DNA transposons, short interspersed nuclear elements (SINEs), long interspersed nuclear elements (LINEs) or Helitrons depending on species, were significantly higher than the whole genome average. This further supports the hypothesis that repetitive elements are the most favorable targets of A-to-I editing in metazoans. Similar results were obtained even if we weighted each editing site with its editing level (Supplementary Fig. 4e,f). Moreover, we observed negative correlations between the divergence rates and the editing densities of repetitive elements in most species (Fig. 3d and Supplementary Fig. 4g), suggesting that A-to-I editing preferentially targets evolutionarily young repetitive elements that likely only relatively recently invaded the genome of each species. Given that hyper-edited dsRNAs can be degraded by endonuclease V ^45^, RNA editing may therefore serve as a guardian mechanism to avoid the overactivation of repetitive elements in metazoans.

### Recoding RNA editing is rare and generally nonadaptive in metazoans

The phenomenon of recoding editing has gained considerable research interest, as it can result in nonsynonymous substitutions in protein-coding sequences, and thus has the potential to increase proteome diversity by introducing novel protein isoforms ^3, 6^. We observed that the number of recoding sites varied greatly across species, with the octopus having an overwhelming higher number (29,464) than all other species (median 850). In general, the proportion of recoding sites among all A-to-I editing sites was low, ranging from less than 1% to 7% in the majority of metazoans, with only 1% to 5% of all expressed protein-coding genes being recoded. However, the proportions of recoding sites in the fruit flies and the sea squirt were prominently high, reaching 33% (711/2,149), 30% (641/2,165) and 14% (850/6,254) in *D. melanogaster*, *D. simulans* and *C. savignyi*, respectively (Fig. 4a). This may possibly be due to the reduced proportion of editing sites in repetitive elements for these species (see Fig. 3a and Supplementary Fig. 4c,d).

**Figure 4.**
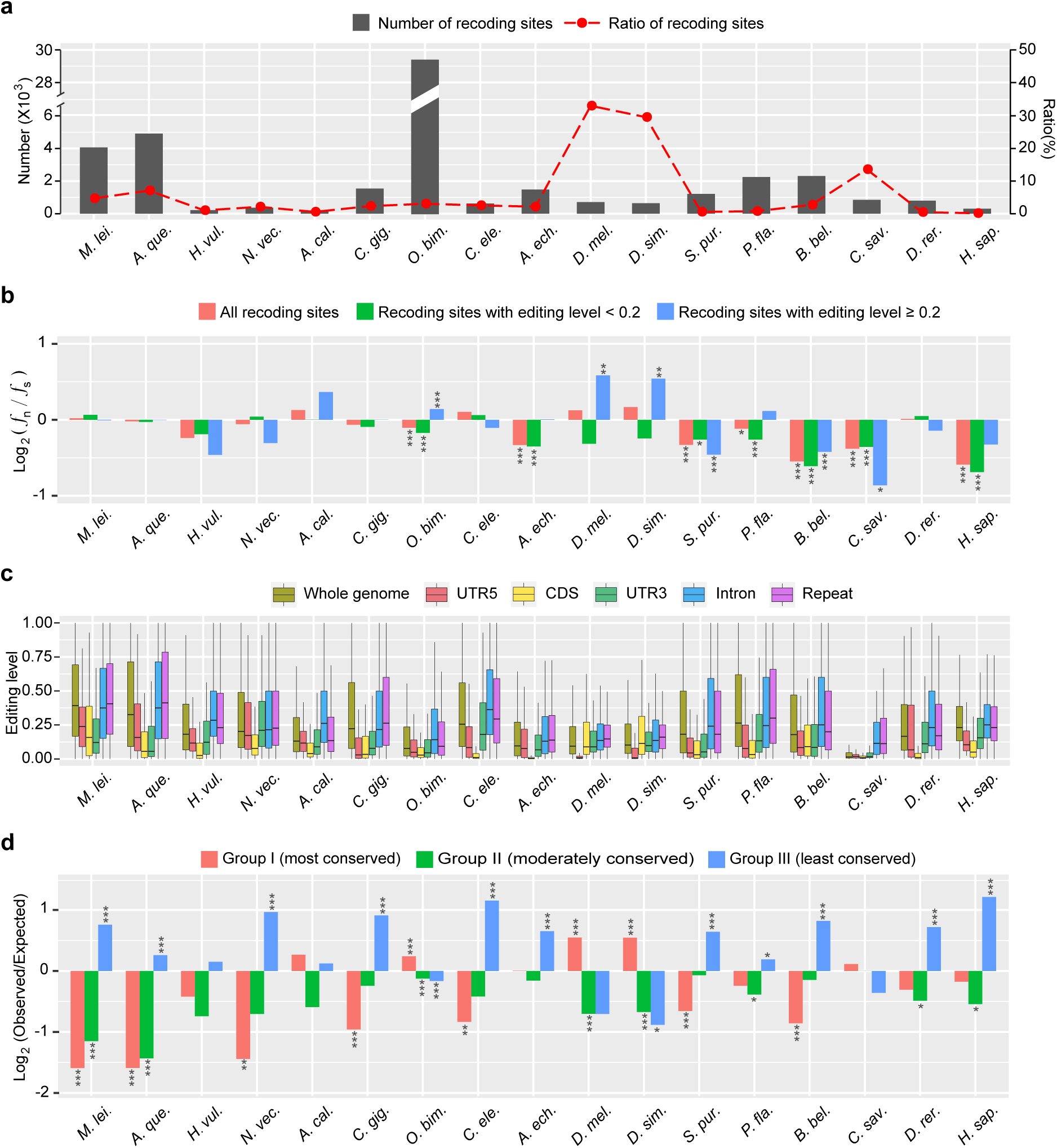
The prevalence and adaptive potential of recoding editing in metazoans. (**a**) The number (left y-axis) and proportion (right y-axis) of recoding sites versus total A-to-I editing sites in each species. (**b**) The frequency of nonsynonymous editing (ƒ_n_) versus synonymous editing (ƒ_s_) for all A-to-I editing sites, and A-to-I editing sites with low (< 0.2) and high (≥ 0.2) editing levels. ƒ_n_ (or ƒ_s_) was calculated as the number of A-to-I editing sites causing nonsynonymous (or synonymous) changes against the number of potential nonsynonymous (or synonymous) adenosine sites if A is replaced with G from the genes with ≥ 1 editing site in their coding regions. Significance levels were estimated by two-tailed Fisher’s exact tests. (**c**) The comparison of editing levels of A-to-I editing sites in different genomic elements. For each editing site in a species, the mean editing level across samples with RNA depth ≥ 10X was calculated and used in this analysis. (**d**) The observed/expected number of genes subject to recoding editing among genes with different conservation levels. The expected probability of a gene being recoded in a species was estimated as the number of recoded genes (i.e. genes with ≥ 1 recoding site) out of all transcribed protein-coding genes (i.e. RPKM > 1 in at least one sample) in this species, and the expected number of recoded genes in each conservation group was calculated as the number of genes in this group multiplied by the expected probability of a gene being recoded. Significance levels were estimated by two-tailed binomial tests. Asterisks in panels **b** and **d** indicate the significance levels with “*” representing *p* < 0.05, “**” < 0.01 and “***” < 0.001.

We next examined the effect of natural selection on recoding sites. It has been previously reported that nonsynonymous editing is generally adaptive in fruit flies and cephalopods ^28–30, 34^. If this is so, one would expect that, in relation to synonymous editing, which is expected to be neutral, the frequency of nonsynonymous editing (ƒ_n_) calculated as the number of A-to-I editing sites causing nonsynonymous changes against all potential nonsynonymous adenosine sites if A is replaced with G, is higher than that of synonymous editing (ƒ_s_) (see Methods). When considering all recoding sites together, we observed that the frequencies of nonsynonymous editing were either close to, or significantly lower than, synonymous editing in all species (Fig. 4b). This therefore argues against the adaptive hypothesis, and suggests that the recoding editing events observed in coding regions of most metazoans are generally neutral or deleterious, consistent with previous reports in humans ^35^. Consistently, editing levels of A-to-I sites in coding regions were generally lower than the genome average and other types of genomic elements (Fig. 4c), implying that editing of coding regions tends to be suppressed. However, when we divided the recoding sites of each species into lowly (editing level < 0.2) and highly (editing level ≥ 0.2) edited groups, we found that the frequencies of nonsynonymous editing in fruit flies and octopus became significantly higher than synonymous editing in the highly edited group (Fig. 4b). This demonstrates a relatively larger portion of adaptive recoding sites exists in these two lineages than in other metazoans.

If recoding editing is generally nonadaptive, one would also expect that nonsynonymous editing is depleted from evolutionarily conserved genes which are less tolerant to mutations. We thus divided the genes of each species into three groups according to the degree of evolutionary conservation (see Methods). Group I and II comprise genes that have orthologs in closely-related species, but with relatively low and high *d_N_*/*d_S_* ratios, representing the most and moderately conserved groups, respectively. Group III comprises all the remaining genes, that cannot find orthologs, and represents the least conserved group. As expected, the genes subjected to recoding editing were generally enriched in the least conserved groups in most metazoans (Fig. 4d), suggesting that recoding editing tends to be purged from the evolutionarily conserved genes in most metazoans. Nevertheless, an inverse tendency can be observed in the fruit flies and octopus, probably due to the relatively larger portions of adaptive recoding sites in these species. This also implies that adaptive recoding editing more likely emerged in the evolutionarily conserved genes, which benefit from increasing protein diversity without introducing DNA mutation in these genes.

### RNA editing preferentially affects cellular communication and cytoskeleton related genes

To uncover the functional preference of genes targeted by A-to-I editing in metazoans, we conducted gene ontology (GO) based functional enrichment analysis for the RNA-recoded genes (i.e. genes with at least one recoding site of which the average editing level across samples > 0.1 or shared by at least two samples) in each species. Consistent with previous observations in mammals ^7, 9^, insects ^28, 29, 31^ and cephalopods ^33, 34^, we found that neurotransmission-related functions such as ion transmembrane transport, synaptic transmission and gated channel activity were significantly enriched in diverse species including human, zebrafish, acorn worm, *Drosophila*, ant and octopus (Fig. 5a), confirming the important role of RNA editing in modulating neural function in bilaterians. Representative examples are the voltage-gated K^+^ channels, that show the same recoding events on two highly conserved amino acid residues within the ion transport domain among *Drosophila*, ant, octopus and even human (Fig. 5b and Supplementary Table 5), and the glutamate ionotropic receptors in vertebrates (Supplementary Fig. 5a and Supplementary Table 5). Interestingly, although a nervous system is absent in the sponge ^46^, functional categories related to cellular communication, signal transduction and response to stimulus were significantly enriched in this early-branching and morphologically simple metazoan. Given that neurotransmission is also part of the cell communication and signal transduction processes which mediate cellular response to internal and external stimulus ^47^, these results imply that RNA editing might have been adopted to modulate the molecular pathways of stimulus response during the early stage of metazoan evolution.

**Figure 5.**
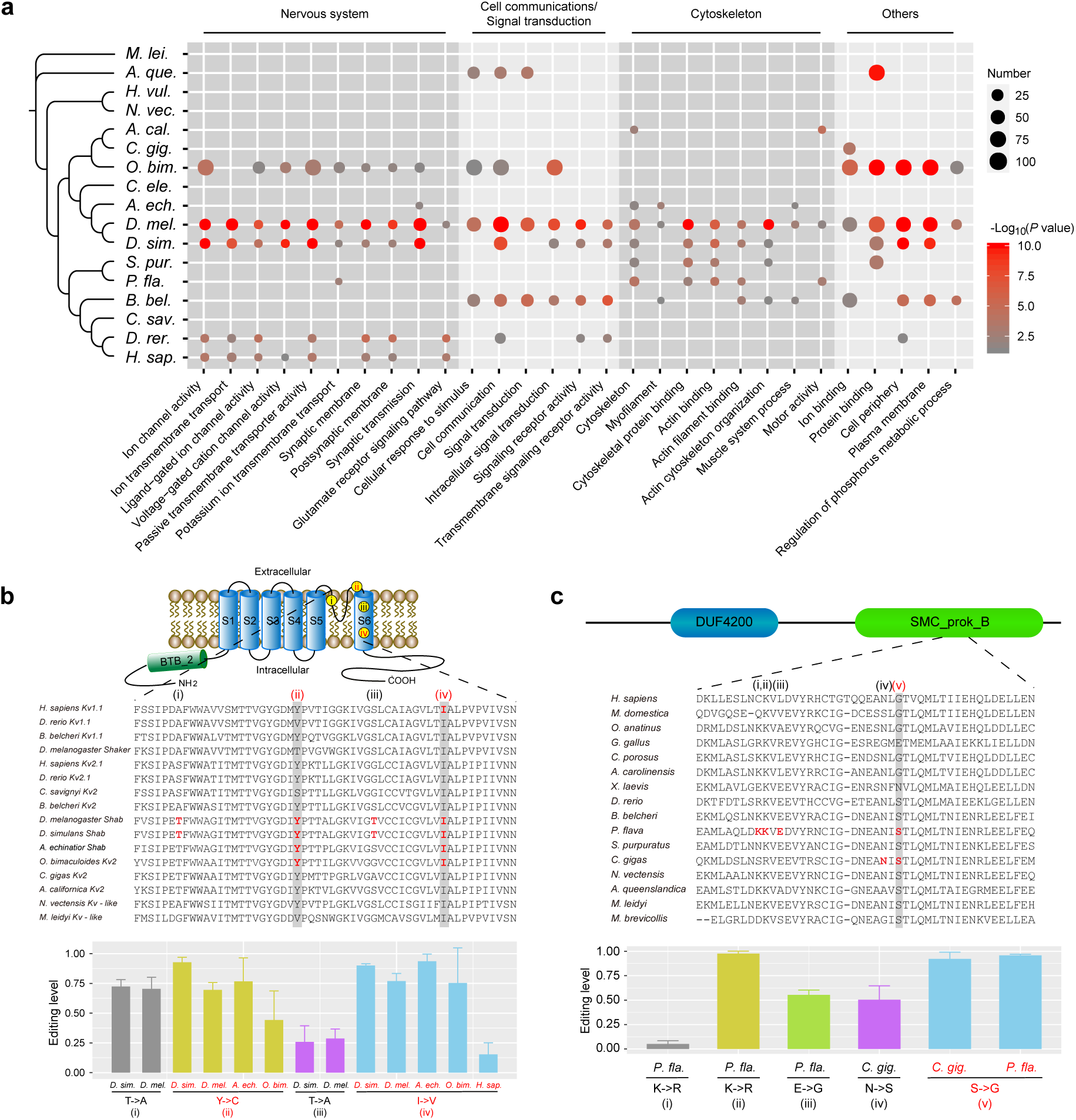
Functional preference of recoding editing in metazoans. (**a**) Functional categories that are enriched by recoded genes in no less than three species (Hypergeometric test adjusted *p* < 0.05). Only genes with at least one recoding site of which the average editing level across samples > 0.1 or shared by at least two samples in each species were used for the functional enrichment analysis (see Methods for details). (**b**) An example showing the convergent evolution of recoding editing in the voltage-gated K^+^ (Kv) channels among *Drosophila*, ant, octopus and human. The upper part shows the classic structure of the Kv channel within the cell membrane that contains six membrane-spanning domains (S1–S6). Yellow dots indicate the locations of recoding sites. The middle part shows the multiple sequence alignment of the region containing the four recoding sites, with recoding sites identified in each species highlighted by red color and the conserved recoding events shared across phyla highlighted by gray shadow. The lower part shows the editing levels of the recoding events in each species. (**c**) An example showing the convergent evolution of a recoding editing event in the same amino acid residue of the cytoskeleton-related gene *CFAP100* between the oyster *C. gigas* and acorn worm *P. flava*. The upper part shows the domain organization of the oyster CFAP100 protein (XP_011420958.1) annotated by the Conserved Domain Database (CDD) of NCBI. The middle and lower part are similar with panel **b**. Error bars in panels **b** and **c** represent s.d. across samples.

However, it is unexpected that significant enrichment of cytoskeleton-related functions such as cytoskeletal protein binding, actin cytoskeleton organization and motor activity, was frequently observed in diverse bilaterians (Fig. 5a). Recoding editing of genes involved in cytoskeleton-related functions has been only rarely reported previously ^33, 34^. The only well-documented cases so far are the actin crosslinking proteins filamin α (*FLNA*) and filamin β (*FLNB*), of which a conserved Q-to-R recoding event occur at the same position of both proteins in mammals ^48^. Our data not only confirms that recoding editing of *FLNA* or *FLNB* occurs in humans, but also detects recoding events in sea urchin (*FLNB*), *Drosophila* (*FLNA*), and acorn worm (*FLNA*). Other representative examples comprise the *cilia and flagella associated protein 100* (*CFAP100*) which contains a S-to-G recoding event shared by oyster and acorn worm (Fig. 5c and Supplementary Table 5), and *fascin* (an actin filament-bundling protein) which has a Q-to-R recoding event shared by octopus, sea urchin and lancelet (Supplementary Fig. 5b and Supplementary Table 5). The repeated emergence of same recoding editing in the cytoskeleton-related genes in different lineages emphasizes an important, but previously unappreciated, role of RNA editing in regulating cytoskeleton-related functions in metazoans.

## Discussion

The phenomenon of RNA editing has been reported previously across a diverse range of eukaryotes including metazoans, protists, fungi and plants, and to affect different types of RNA^1, 49^. However, while in most eukaryotes it is exclusively limited to mitochondrial or chloroplast RNA, the extensive editing of nuclear-transcribed mRNA is phylogenetically rare, and restricted to metazoans and some filamentous ascomycetes in which it originated through independent mechanisms ^3, 50^. In this study, we present the first direct evidence that this method for extensive alteration of nuclear DNA-encoded genetic information was adopted alongside the origin of *ADARs* by the last common ancestor of extant metazoans ca 800 million years ago ^51^, following its divergence from unicellular choanoflagellates. This raises the possibility that ADAR-meditated RNA editing is an ancient regulatory process that was fundamental for initial metazoan evolution. The evolutionary maintenance of ADAR-meditated RNA editing in almost all extant metazoan lineages also emphasizes its essentiality in metazoan biology.

Consistent with the evolutionary constraint of ADARs, we show that the nucleotide sequence and structural features surrounding A-to-I editing sites, including the strong favor of uridine/adenosine and disfavor of guanosine in the adjacent 5’ positions, and the tendency of the underlying sequences to form dsRNA structures, are under strong constraint across the animal kingdom, from the earliest branching ctenophore and sponge to human. These findings might be valuable for ADAR-based RNA engineering, such as the recently reported RESTORE and LEAPER approaches, which can recruit endogenous ADAR to specific transcripts for site-directed RNA editing in human cells ^52, 53^, as these conserved features imply that the approaches developed based on one species (usually human) may well be easily applicable to other metazoan species with *ADARs*.

It is now generally acknowledged that the complexity of transcriptional regulation coincides with organismal complexity ^54^. RNA editing and alternative splicing have long been proposed to serve as important co/post-transcriptional regulatory mechanisms for increasing transcriptome diversity ^3, 55^. However, while alternative splicing has been demonstrated to be strongly associated with organismal complexity ^56^, we do not observe such a relationship between the extent of global RNA-editing and organismal complexity in metazoans. Together with our observations that in metazoans A-to-I editing preferentially targets evolutionary young repetitive elements, and that recoding events in protein-coding sequences are generally neutral or slightly deleterious, these findings question the ancestral role of A-to-I RNA editing as a transcriptome or proteome diversifier in metazoans. Recent *ADAR1*-knockout studies in human cells and mice indicated that ADAR1-mediated A-to-I editing of endogenous dsRNAs formed by inverted repeats, plays a key role in preventing cellular sensing of endogenous dsRNA as nonself (e.g. viral RNA), thus avoiding autoinflammation ^18, 57^. This suggests that the avoidance of aberrant immune responses triggered by the accumulation of endogenous dsRNA represents the primary driving force for preserving the extensive A-to-I editing in most metazoan lineages. Alternatively, given that most editing sites are only edited at low to moderate levels in all the species examined, and thus might not be sufficient to unwind dsRNAs to avoid immune response, we hypothesize that metazoans may benefit from the maintenance of mild single-nucleotide mutations in the RNA pool, as these mutations can provide plentiful transcript variants that might help metazoans cope with unpredictable future conditions in their life.

Our extensive survey across the phylogeny of metazoans also emphasizes that *Drosophila* and cephalopods, whose RNA editomes harbor relatively high proportions of adaptive recoding events subject to positive selection, are actually evolutionary exceptions in the animal kingdom. The abundant recoding editing in cephalopods has been demonstrated to emerge in the ancestor of coleoids after splitting from nautiloids, with the expansion of the cephalopod RNA editomes^34^. In contrast, we find that the *Drosophila* RNA editomes have been greatly contracted in comparison to most metazoans, while a considerable portion of recoding events is maintained by natural selection. When this *Drosophila* pattern emerged during the evolution of insects remains unknown. At least, the fact that more ‘classic’ RNA editomes, in which the majority of sites targeting repetitive elements and rare recoding editing, are observed in ants and recently in bumble bees ^32^, indicates that this *Drosophila* pattern must emerge after the divergence of Diptera and Hymenoptera ca 345 million years ago ^58^.

RNA editing has been long acknowledged to regulate neural functions, affecting genes encoding ion channels and neuroreceptors ^7, 9, 10^, consistent with the results of our functional enrichment analysis of recoded genes in diverse species. Thus what is most surprising about our observations is the over-representative of recoded genes encoding cytoskeleton-related functions in diverse species, implying that post-transcriptional diversification of cytoskeleton-related genes via RNA editing might be an important way through which to increase cellular complexity during the evolution of metazoans. In particular, in some cases, we find exactly the same positions are edited and cause the same amino acid changes in evolutionarily conserved residues in distantly related species. The cytoskeleton is an interconnected network of filamentous polymers and regulatory proteins, which carries out broad functions including spatially organizing the contents of the cell, connecting the cell physically and biochemically to the external environment and generating coordinated forces that enable the cell to move and change shape ^59^. It will be necessary for future studies to ascertain which aspect of cytoskeleton-related functions RNA editing regulate.

In summary, our study provides the first large-scale and unbiased transcriptome-wide investigation of RNA editing across the phylogeny of metazoans. These resources are valuable for our understanding of the biological role and evolutionary principle of RNA editing in the animal kingdom.

## Methods

### Sample collection

To rule out that false positives resulted from genetic variation during RNA-editing site identification, matching DNA and RNA sequences generated from the same individual/specimen are the ideal data for use in RNA editing studies ^39, 60^. Thus, for the metazoan species with sufficient body mass, both genomic DNA and total RNA were extracted from the same individual, after grinding of the tissue/whole organism in liquid nitrogen. Two to three individuals were collected as biological replicates. These species included the comb jelly *Mnemiopsis leidyi* (three whole adults), the sponge *Amphimedon queenslandica* (three biopsies from three adults), the sea anemone *Nematostella vectensis* (three whole adults), the sea hare *Aplysia californica* (three whole juveniles), the oyster *Crassostrea gigas* (three whole adults after removing shells), the sea urchin *Strongylocentrotus purpuratus* (three pairs of gonad and non-gonad tissues dissected from one female and two male adults; non-gonad tissues comprised the digestive, water vascular, and nervous systems), the acorn worm *Ptychodera flava* (three whole adults), the lancelet *Branchiostoma belcheri* (three whole adults), the sea squirt *Ciona savignyi* (two whole adults) and the zebrafish *Danio rerio* (three whole adults).

For metazoan species from which a single individual is not sufficient to allow the simultaneous extraction of sufficient DNA and RNA for sequencing library construction, 10-15 individuals with similar genetic background were pooled together, then both genomic DNA and total RNA were extracted from the same pool of organisms after the whole pool was ground in liquid nitrogen. These included the hydra *Hydra vulgaris* (10 adults per pool, two pools to serve as biological replicates), the fruit fly *Drosophila melanogaster* (15 male adults per pool, two pools), and *Drosophila simulans* (15 male adults per pool, two pools).

For the unicellular species and tiny metazoan species, biomass was first increased by the propagation of a single colony with the same genetic background, then both genomic DNA and total RNA were extracted from the same culture of organisms. These included the ichthyosporean *Sphaeroforma arctica* (three cultures to serve as biological replicates), the filasterean *Capsaspora owczarzaki* (three cultures), the choanoflagellate *Salpingoeca rosetta* (three cultures) and *Monosiga brevicollis* (three cultures), and the metazoan *Trichoplax adhaerens* (three cultures).

All the species were either collected from conventionally grown lab conditions, or obtained from the wild. With the exception of the sea hare samples which were purchased from the National Resource for Aplysia, University of Miami, 4600 Rickenbacker Causeway, Miami, FL 33149, samples of all the other species were kindly provided by researchers who have worked on corresponding species for years. The strain identifier (if applicable), geographical origin and providers of each species were listed in Supplementary Table 1.

Genomic DNA of all species was extracted with the phenol/chloroform/isopentanol (25:24:1) protocol. The integrity of the DNA samples was assayed by agarose gel electrophoresis (concentration: 1 %; voltage: 150 V; Time: 40 min) before DNA-seq library construction. Total RNA of all species except the choanoflagellates was extracted using TRIzol Reagent according to manufacturer’s protocol (Invitrogen, CA, USA). Total RNA of the choanoflagellates *S. rosetta* and *M. brevicollis* was extracted using the RNAqueous Kit (Ambion, CA, USA). The quality of the RNA samples was assayed by the Agilent 2100 Bioanalyzer (Thermo Fisher Scientific, MA, USA) before RNA-seq library construction. In summary, a total of 53 DNA and 53 RNA samples were obtained in this study. After quality control before library construction, two out of the three RNA samples of *M. brevicollis* and one out of the three RNA samples of *N. vectensis* were discarded due to poor RNA integrity (RIN < 6).

### Library construction and sequencing

The strand-specific RNA-seq libraries for all the RNA samples were prepared using the TruSeq Stranded mRNA LT Sample Prep kit (RS-122-2101, Illumina) with 1 µg total RNA as input, then sequenced on the Illumina HiSeq 4000 platform using the PE100 chemistry, according to the manufacturer’s instructions (Illumina, San Diego, CA, USA).

The genomic DNA samples were either sequenced on an Illumina HiSeq 4000 or a BGISEQ-500RS platform. The Illumina HiSeq and BGISEQ-500 platforms have been proved to generate data with comparable quality and show high concordance for calling single nucleotide variants by multiple independent studies ^61–63^. For the Illumina DNA libraries, 1 μg genomic DNA per sample was fragmented by a Covaris ultrasonicator, followed by end repair, 3′-end addition of dATP and adapter ligation. The ligated fragments were then size selected at 300 bp on an agarose gel and amplified by 10 cycles of PCR. The amplified libraries were purified using the AxyPrep Mag PCR Clean-Up Kit (Axygen, MA, USA) then sequenced on the Illumina HiSeq 4000 platform using the PE100 chemistry according to the manufacturer’s instructions (Illumina, San Diego, CA, USA). The BGISEQ DNA sequencing libraries were prepared using the MGIEasy DNA Library Prep Kit (V1.1, MGI Tech) with 1 μg genomic DNA as input, and sequenced on the BGISEQ-500RS platform using the PE100 chemistry according to the manufacturer’s instructions (MGI Tech Co., Ltd., Shenzhen, China). Details about the sequencing platform and data production for each sample were presented in Supplementary Table 1.

### Identification of RNA-editing sites

#### (i) Quality control for raw sequencing data

All the DNA- and RNA-seq reads were first submitted to SOAPnuke (v1.5.6) ^64^ for quality control by removal of adapter-contaminated reads and low-quality reads before subsequent analyses with parameters *-G -l 20 -q 0.2 -E 60 -5 1 -Q 2*.

#### (ii) Adjustment of reference genome with DNA-seq data

Given that many samples were collected from wild animals, which have high levels of heterozygosity, or were from strains which are genetically different from those used for assembling the reference genomes, we employed Pilon (v1.21) ^65^ to adjust the reference genome of each species using the DNA-seq data from different samples separately, generating sample-specific reference genomes for each species before RNA-editing site identification. Specifically, DNA sequence reads from each sample of a species were first aligned to the published reference genome using BWA-MEM (v0.7.15) ^66^ with default parameters. Then, genome adjustment was performed by Pilon with default parameters except that *--fix snps* was set, using the original reference genome FASTA and the DNA BAM files as input. It is noteworthy that we only adjusted SNPs in the reference genomes in order to ensure that the adjusted genomes from different samples of the same species have the same length and the same coordinate system. The version and source of the original reference genome for each species were listed in Supplementary Table 1.

#### (iii) Identification of RNA-editing sites with RES-Scanner

RNA-editing sites from each sample were first identified by RES-Scanner (v20160713), a software package that was designed to identify transcriptome-wide RNA-editing sites with matching DNA- and RNA-seq data from the same individual or specimen ^39^. Briefly, RES-Scanner invoked BWA-ALN (v0.7.15) ^67^ to align the DNA and RNA reads that passed quality control to the adjusted reference genome of each species, followed by filtering low-quality alignments, calling homozygous genotype from DNA data, and identifying candidate RNA-editing sites from RNA data by ruling out false-positives resulted from genetic variants and sequencing or alignment errors. In general, default parameters were used for the whole pipeline, except that the mapping quality cutoff was set to 5 for DNA alignment (default 20) and the numbers of bases masked at the 5’- and 3’-end of a DNA read was set to 0 (default 6). This was done as we found that lowering these requirements for the DNA data could yield RNA-editing sites with higher accuracy in many species, manifesting as the higher proportions of A-to-I editing sites out of all identified editing sites.

#### (iv) Identification of hyper-editing sites

Given that A-to-I editing sites tend to occur in clusters, the heavily edited RNA reads (commonly called hyper-edited reads) which contain many of the same type of substitutions in relation to the reference genome, often fail to be aligned during normal alignment process. In order to capture these hyper-edited reads and the clusters of editing sites they harbor, we next performed hyper-editing detection for each sample following a scheme originally proposed by Porath *et al* ^40^.

We first collected the RNA read pairs that could not be aligned to the adjusted reference genome or that had mapping quality < 20 from the RNA BAM files generated by the RES-Scanner pipeline as described above. We then removed the read pairs for which one or both reads contained more than 10% of Ns along their lengths, or had particularly large (>60%) or small (<10%) percentage of a single-type nucleotide as recommended by Porath *et al* ^40^. Next, we adopted a “three-letter” alignment strategy to align these potential hyper-edited reads, in order to overcome the excess mismatches in relation to the reference genome. For example, to align the RNA reads with many A-to-I editing sites (i.e. many A-to-G mismatches), all Ts in the first read of a read pair were transformed to Cs, and all the As in the second read of a read pair were transformed to Gs. This is because, for read pairs generated from the dUTP-based strand-specific RNA-seq libraries, the second read is from the original RNA strand/template while the first read is from the opposite strand ^68^. In the meantime, two versions of the reference genome were created, of which the first version was named the *positive* reference, with all As transformed to Gs, and the second version was named the *negative* reference, with all Ts transformed to Cs.

Next, the transformed read pairs were aligned to both the *positive* and *negative* references by BWA-ALN with parameters *-n 0.02 -o 0*, yielding the *positive* and *negative* alignments, respectively. Then, we filtered both alignments by removing read pairs that were not aligned to the reference genome concordantly, and the reads within concordantly aligned pairs that had mapping score < 20. In addition, for *positive* alignment, we further required that the first read in a pair was the reverse complement of the reference genome, while the second read was aligned to reference genome directly; for *negative* alignment, we required that the first read in a read pair was directly aligned to reference genome, while the second read was the reverse complement of the reference genome.

After the strict quality control for the BWA alignments, we converted the transformed reads to their original sequences, followed by trimming the first and last 10 bases of each read in the alignments. Then we identified hyper-edited reads by requiring the mismatch rate of a trimmed read to be > 5%, and the proportion of the expected mismatches (i.e. A-to-G substitution in this example) against all mismatches to be > 60% as recommended by Porath *et al* ^40^. Finally, BAM files of hyper-edited RNA reads were submitted to RES-Scanner to extract potential editing sites together with the matching DNA BMA files generated in the previous step. RES-Scanner was run with default parameters in general, except that the mapping quality cutoff was set to 5 for DNA alignment, the numbers of bases masked at the 5’- and 3’-end of a read were set to 0 for both DNA and RNA reads, the minimum number of RNA reads supporting editing was set to 2 (default 3), and the minimum editing level was set to 0 (default 0.05).

The above hyper-editing detection method was undertaken for all of the 12 possible substitution types of RNA editing in each sample of a species, and the results from all the 12 substitution types were combined together by discarding those sites that presented different editing types in any single genomic position.

#### (v) Combing the results of RES-Scanner and hyper-editing detection

To generate the representative RNA-editing sites for a species, and to improve the identification of editing sites in each sample, we combined the editing sites identified by RES-Scanner (step iii) and hyper-editing detection (step vi) in each sample, to obtain a comprehensive map of potentially editable positions in the reference genome of each species. If a genomic position was identified as an editing site in both methods, we respectively added the numbers of RNA reads supporting editing, and the number supporting non-editing as generated by these two methods. We then retrieved the missed editing sites in each sample in these editable positions using the criteria of at least one RNA read supporting editing and the false discovery rate (FDR) ^69^ adjusted *p* value for this site to be resulted from sequencing error < 0.01. Specifically, statistical tests were performed based on the binomial distribution B(*k*, *n*, *p*), where *p* was set to be the maximal probability of an RNA base to be a sequencing error (i.e. 0.1% here as we only used RNA bases with Phred quality score ≥ 30), *n* was equal to the total read depth of a given candidate editing site, and *k* denoted the number of reads supporting editing. We also used the DNA-seq data from multiple samples to further remove false-positives resulted from genetic variants, by discarding those editing sites for which the genomic DNA showing the same type of substitution as RNA editing (i.e. the frequency of edited base versus the total number of bases covering this position > 0.1) in any one of the multiple DNA samples. RNA-editing sites that displayed different editing types in different samples of a species were also discarded.

We have updated the software package RES-Scanner we previously established for RNA-editing site scanning by compiling above steps (step *i* to *v*). This RES-Scanner2 now can also identify hyper-editing sites. It works from raw sequencing reads and is applicable to RNA-editing site detection in any species with matching DNA- and RNA-seq data.

### RNA-editing sites for additional metazoan species

To increase the phylogenetic coverage of the investigated species, we collected the matching DNA-seq and strand-specific RNA-seq data from the nematode *Caenorhabditis elegans* (pooled whole organisms collected from three larval stages and two adult stages) ^26^, the leaf-cutting ant *Acromyrmex echinatior* (three pooled head samples of the small worker caste collected from three colonies, respectively) ^31^, the octopus *Octopus bimaculoides* (four neural tissue samples including faxial nerve cord, optic lobe, subesophageal ganglia and supraesophageal ganglia) ^37^ and human (three brain samples from three male adults, respectively) ^22^. The SRA accession numbers and statistics of the downloaded sequencing data were presented in Supplementary Table 1. RNA-editing sites in each of the four species were identified using the same procedure (step *i* to *v*) as described above.

### Refining the ORFs and annotating UTRs for protein-coding genes

Protein-coding genes (GFF/GTF and corresponding cds/pep FASTA files) were downloaded from public databases along with the reference genomes, of which the sources were presented in Supplementary Table 1. The correctness of the open-reading frames (ORFs) in the GFF/GTF files were checked for all the protein-coding genes, with the defective ORFs such as those that were not the integer multiple of 3 in length or not exactly matching the protein sequences presented in the downloaded pep FASTA files being carefully corrected by in-house scripts. Then the transcript model with the longest ORF was chosen as the representative model for a locus if multiple transcript models were annotated in this locus.

5’- and 3’-UTRs for the representative ORFs were annotated using the RNA-seq data used in this study, for all the species except for human. Briefly, RNA-seq reads that passed quality control as described above were first aligned to the reference genome of each species by HISAT2 (v2.1.0) ^70^, with default parameters except setting *--rf*, followed by removing those reads that could be mapped to multiple positions of the genome. Then, transcribed regions with continual RNA depth ≥ 5X were extended from the 5’- and 3’-end of each representative ORF to serve as initial 5’- and 3’-UTRs, respectively. Next, an iterative process was used to further recruit the upstream or downstream transcribed regions that were apart from, but linked by ≥ 5 junction reads to previously defined UTRs. If a gene had different 5’- or 3’-UTRs annotated in different samples, the longest one was chosen as the representative 5’- or 3’-UTR for this gene.

### Gene expression quantification and transcript assembly with RNA-seq data

HISAT2 alignments generated in the above analysis were used to quantify gene expression levels for the refined representative gene models in each species. Only the RNA-seq reads that were aligned to one position of the reference genome, and that overlapped with annotated exons were kept for expression quantification. Gene expression levels were measured by RPKM (reads per kilobase per million mapped exonic reads), and the RPKM values in all the sequenced samples from the same species were adjusted by a scaling normalization method based on TMM (trimmed mean of M values) to normalize the sequencing bias among samples^71^. We also assembled transcripts for each species with StringTie (v1.3.4d) ^72^ with default parameters using the HISAT2 alignments as input. These transcript models were regarded as one kind of reference models during the manual annotation of *ADAR* genes as described below.

### Annotation of *ADAR* genes in each species

ADAR protein sequences of *Nematostella vectensis* (XP_001642062.1 and XP_001629615.1), *Drosophila melanogaster* (NP_569940.2), *Caenorhabditis elegans* (NP_492153.2 and NP_498594.1), *Crassostrea gigas* (EKC20855.1 and EKC32699.1), *Strongylocentrotus purpuratus* (XP_011680614.1 and XP_781832.1), *Ciona intestinalis* (XP_002128212.1), *Danio rerio* (NP_571671.2, NP_571685.2, XP_021334693.1 and XP_686426.5) and *Homo sapiens* (XP_024305442.1, NP_056648.1 and NP_061172.1) collected from NCBI were used as queries to search for *ADAR* genes in reference genomes of all the 22 species by TBLASTN (blast-2.2.23) ^73^ with parameters *-F F -e 1e-5*, followed by the determination of gene structure and protein sequences in the target species with GeneWise (wise2.2.0) ^74^. The predicted proteins were then aligned to the NCBI nr database to confirm whether they were ADARs.

Next, we manually compared the gene models in the putative *ADAR* loci resulted from homologous predictions, transcript assemblies by StringTie and the published gene set of each species, and we chose the models with the longest ORFs as the representative models. Domain organizations of the manually confirmed ADAR proteins were predicted using the CD-Search tool in NCBI (CDD v3.17; https://www.ncbi.nlm.nih.gov/Structure/cdd/wrpsb.cgi) and Pfam (release-32.0; https://pfam.xfam.org) with default settings, and only ADARs with at least one dsRNA binding domain (dsRBDs) and one adenosine-deaminase domain (AD) were regarded as potential *ADAR* genes. Of note, *ADAD* genes, which usually contain one or more dsRBDs and one AD, were also identified as potential *ADARs* by our criteria, but they could be distinguished from *ADARs* according to phylogenetic analysis (see below). The information of *ADAR* genes annotated in each species, including the coding nucleotide sequences, protein sequences, domain annotations and editing sites are presented in Supplementary Table 3.

Phylogenetic analysis of all the potential ADARs identified above, were performed with the AD peptide sequences (ca 324 amino acids in length) using MEGA7 with the neighbor-joining method ^75^. We did not perform phylogenetic analysis whit the dsRBDs, as the lengths of the dsRBDs were generally very short (ca 40 to 60 amino acids) and the copy number of dsRBDs varied among ADARs both within and between species. The peptide sequences of ADs used for phylogenetic analysis were aligned using ClustalW as implemented in MEGA7. Reliability of the trees was estimated using 1,000 bootstrap replications (Supplementary Fig. 2d). To further estimate the divergence between any two potential ADARs, we calculated the nonsynonymous substitution rates (*d_N_*) for any pair of potential ADARs using PAML (v4.9i) ^76^ with the Yang & Nielsen (2000) method ^77^, according to the codon alignment of the ADs (Supplementary Fig. 2e).

### Identification of editing sites locating in potential dsRNA regions

The dsRNA regions formed by two potential mechanisms, intramolecular folding of pre-mRNA and intermolecular hybridization of sense-antisense transcripts, were tested for the enrichment of A-to-I editing sites.

For the mechanism of intramolecular folding, we extracted a 401 nt sequence centered on each A-to-I editing site, then searched this query sequence against a 4001 nt sequence centered on corresponding A-to-I editing site using BLASTN (v2.2.26) with parameters *-F F -e 1e-2*. Then an A-to-I editing site was identified as locating in a dsRNA region formed by intramolecular folding, if a reverse-complement alignment was detected with identity ≥ 80%, the aligned length was ≥ 50 nt, and the aligned region of the query sequence spanned the edited adenosine. For the mechanism of intermolecular hybridization of sense-antisense transcripts, we examined the RNA coverage of a 101 nt region centered on each A-to-I editing site, and searched for the regions with RNA depth ≥ 2X along >50% of the region length, on both strands.

To estimate the expected ratio of A-to-I editing sites that occurred in dsRNA regions formed by the above two different mechanisms in each sample, we randomly selected an adenosine site with comparable RNA depth (i.e. within ± 20% of the editing site) for each editing site in a sample, and performed the same analyses for these control adenosine sites. The significance levels for the difference between the observed and expected ratios were examined by two-tailed paired t-tests in each species.

### Definition of clustered and isolated editing sites

For each sample of a species, we considered a genomic region containing ≥ 3 A-to-I editing sites, of which the distance for two adjacent sites was ≤ 30 nt, as an RNA-editing cluster. The genomic locations of the first and last editing sites in a cluster were assigned as the start and end genomic positions of this cluster. A-to-I editing sites located in the defined editing clusters were regarded as clustered editing sites, and those outside editing clusters were regarded as isolated editing sites. To estimate the expected ratio of A-to-I editing sites occurring in clusters in each sample, we randomly selected an adenosine site with comparable RNA depth (i.e. within ± 20% of the editing site) for each editing site in a sample, and calculated the ratio of these control adenosine sites occurring in clusters. The significance levels for the difference between the observed and expected ratios were examined by two-tailed paired t-tests in each species.

### Analysis of the neighboring nucleotide preference for A-to-I editing

The Two Sample Logo software (v1.21) ^78^ was used to analyze the neighboring nucleotide preference of A-to-I editing sites with parameters *-K N -T binomial -C nucleo_weblogo -y*. Specifically, for each species, the eleven-nucleotide sequences with the edited adenosines in the center were used as the foreground dataset, while the eleven-nucleotide sequences centered by the transcribed (RNA depth ≥ 2X) but unedited adenosines locating within ± 50 nt of the edited adenosines, were used as the background dataset for Two Sample Logo analysis. Nucleotides were plotted using the size of the nucleotide that was proportional to the difference between the foreground and background datasets.

### Annotation of repetitive elements

Considering that the repetitive elements of many species investigated in this study are either not well annotated and/or not publicly available, we re-annotated the repetitive elements of all the sampled species except human using the same strategy. Repetitive elements of the human genome (GRCh38/hg38) have been well annotated and thus were downloaded from UCSC directly.

Repetitive elements in the genome assembly of other sampled species were identified by homology searches against known repeat databases and *de novo* predictions as previously described ^79^. Briefly, we carried out homology searches for known repetitive elements in each genome assembly by screening the Repbase-derived RepeatMasker libraries with RepeatMasker (v4.0.6; setting *-nolow -no_is -norna -engine ncbi*) ^80^ and the transposable element protein database with RepeatProteinMask (an application within the RepeatMasker package; setting *-noLowSimple -pvalue 0.0001 -engine ncbi*). For *de novo* prediction, RepeatModeler (v1.0.8) ^81^ was executed on the genome assembly to build a *de novo* repeat library for each species, respectively. Then RepeatMasker was employed to align the genome sequences to the *de novo* library for identifying repetitive elements. We also searched each genome assembly for tandem repeats using Tandem Repeats Finder (v4.07) ^82^ with parameters *Match=2 Mismatch=7 Delta=7 PM=80 PI=10 Minscore=50 MaxPeriod=2000*. To confirm the reliability of our annotations, we compared our repeat annotation results of the fruit fly *Drosophila melanogaster* and the zebrafish *Danio rerio* with those downloaded from UCSC and observed good consistency (Supplementary Fig. 3a,b).

### Calculation of RNA-editing density for different genomic elements

To compare the probability of different genomic elements targeted by A-to-I editing, including the protein-coding genes related elements (5’-UTR, CDS, intron and 3’-UTR) and the repeat-associated elements (SINE, LINE, LTR, DNA transposon, Helitron, tandem repeat and other unclassified repeat loci), we calculated the A-to-I editing density for each type of genomic element by counting the number of A-to-I editing sites located in this element type, out of the total number of transcribed adenosines (RNA depth ≥ 2X) from this element type. The editing density of each element type was first calculated for each sample of a species separately, then the mean editing density across samples was calculated as the representative value for a species.

When calculating the editing-level-weighted editing densities for each element type, an editing site with for example an editing level of 0.1, would be regarded as 0.1 editing site instead of 1 editing site, when counting the number of editing sites for an element type. Only editing sites and transcribed adenosines with RNA depth ≥ 10X were used in the weighted analysis.

### Analysis of relationship between repeat divergence and editing density

The divergence rates of repetitive elements in each species were estimated by RepeatMasker, by comparing the repeat sequences to the ancestral consensus sequences identified by RepeatModeler during the repeat annotation process as described above. Only the transcribed repeat loci with no less than 50 nucleotides covered by ≥ 2 RNA reads were used for this analysis. The transcribed repeat loci were first sorted according to divergence rate from the lowest to the highest (i.e. the youngest to oldest), then divided into 10 equal bins with the same transcribed repeat loci in each bin. Next the editing density for each bin was calculated, as the number of A-to-I editing sites located in repeat loci belonging to this bin, divided by the total number of transcribed adenosines (RNA depth ≥ 2X) from the repeat loci in this bin. The editing density of each bin was first calculated for each sample of a species separately, then the mean editing density across samples was calculated as the representative value for a species. The relationships between repeat divergence rate and editing density in all species were displayed by a heatmap as presented in Fig. 3d.

### Estimating the potentials of repeat and non-repeat regions to form dsRNA

The potential of repeat and non-repeat genomic regions to form dsRNA was approximatively measured as the ratios of repeat and non-repeat derived genomic sites locating in regions that could find a reverse-complement alignment in nearby regions. Briefly, we randomly selected 100,000 sites from the genomic regions annotated as repeat and non-repeat, respectively. Then, we extracted a 401 nt sequence centered on each randomly selected site and searched this query sequence against a 4001 nt sequence centered on the corresponding repeat or non-repeat genomic site using BLASTN (v2.2.26) with parameters *-F F -e 1e-2*. Then a repeat or non-repeat derived genomic site was regarded as locating in a potential dsRNA region formed by intramolecular folding, if a reverse-complement alignment was detected with identity ≥ 80%, aligned length ≥ 50 nt, and the aligned region of the query sequence spanned this randomly selected site. The ratio of such sites against all randomly selected sites was calculated to represent the potential of repeat or non-repeat regions to form dsRNA in a species, and the same process was iterated for 100 times to estimate the distribution (see Supplementary Fig. 4c).

### Analyzing the adaptive potential of recoding editing

Recoding editing sites were identified as the sites where the editing events could cause nonsynonymous changes in protein-coding regions. Given that the numbers of recoding sites were generally small in most species, for the evolutionary analysis of recoding editing, recoding sites from different samples of a species were first combined according to their genomic locations. The editing level of a combined recoding site was measured as the mean editing level across samples with RNA coverage ≥ 10X in this position.

To examine the adaptive potential of recoding editing in a species, we compared the frequency of nonsynonymous editing (ƒ_n_) to the frequency of nonsynonymous editing (ƒ_s_) as previously described ^35^. Specifically, ƒ_n_ was calculated as the number of A-to-I editing sites causing nonsynonymous changes (*n*), divided by the number of potential nonsynonymous adenosine sites (RNA depth ≥ 2X in at least one sample) if A is replaced with G (*N*) from the genes with ≥ 1 editing site in their coding regions. ƒ_s_ was calculated as the number of A-to-I editing sites causing synonymous changes (*s*), divided by the number of potential synonymous adenosine sites (RNA depth ≥ 2X in at least one sample) if A is replaced with G (*S*) from the same set of genes. If recoding editing is generally adaptive in a species, one would expect that ƒ_n_ is significantly larger than ƒ_s_ in this species. The significance level for the difference between ƒ_n_ and ƒ_s_ in a species was assessed by a two-tailed Fisher’s exact test using the values of *n*, *N*, *s* and *S* from this species.

To compare the adaptive potential for recoding sites with different editing levels, the same analyses were performed for recoding sites with relatively high (≥ 0.2) and low (< 0.2) editing levels separately, using the sites with RNA depth ≥ 10X and the genes with one or more editing sites achieving this RNA depth in their coding regions.

### Analyzing the evolutionary conservation of recoded genes

Recoded genes were defined as the protein-coding genes with at least one recoding site. To evaluate the evolutionary conservation of the recoded genes in the seventeen species with reliable A-to-I editing (the target species), we identified the orthologous gene of each recoded gene in a closely-related species with a publicly available reference genome (the related species), and calculated the *d_N_/d_S_* ratio (i.e. the ratio of the number of nonsynonymous substitutions per nonsynonymous site (*d_N_*) to the number of synonymous substitutions per synonymous site (*d_S_*)) for each orthologous pair. The closely-related species chosen for each target species is presented in Supplementary Table 6.

Briefly, all the protein sequences from each target species were first aligned to its related species genome using TBLASTN (blast-2.2.26) with parameters *-F F -e 1e-5*, followed by chaining the syntenic blocks and picking one candidate locus for each target-species protein with the highest TBLASTN bit score by in-house scripts. Then the genomic sequences of these candidate loci together with 2 kb flanking sequences, were extracted from the related-species genome and submitted to GeneWise (wise-2.4.1) to determine the protein sequences by aligning the target-species proteins to these related-species genomic sequences. The related-species proteins were then aligned back to all the protein sequences of the target species using BLASTP (blast-2.2.26) with parameters *-F F -e 1e-5*, and only those hitting the expected proteins in the target species with the highest BLASTP bit score were identified as orthologous proteins in related species. Next, the protein sequences of each orthologous pair were aligned using MAFFT (v6.923) ^83^ with parameters *--maxiterate 1000 –localpair*, followed by the replacement of the amino acids by their corresponding codons for each species. The orthologous pairs of which the MAFFT alignments with invalid sites (i.e. presented as “-” in one of the two aligned sequences) exceeding 50% of the alignment length were discarded. Then the *d_N_/d_S_* ratio for each qualified orthologous pair was calculated using PAML (v4.9i) ^76^ with the Yang & Nielsen (2000) method ^77^.

Finally, the genes of each target species were divided into three groups according to the degree of evolutionary conservation, and the observed/expected number of recoded genes among different groups was calculated. Specifically, group I was comprised of genes with orthologs in closely-related species and *d_N_*/*d_S_* ratios lower than the median value among all orthologous pairs, representing the most conserved group; Group II was comprised of genes with orthologs in closely-related species with *d_N_*/*d_S_* ratios higher than the median value among all orthologous pairs, representing the moderately conserved group; Group III was comprised of all the remaining genes that cannot find orthologs in closely-related species, representing the least conserved group. The expected probability of a gene being recoded in a species was estimated as the number of recoded genes out of all transcribed protein-coding genes (RPKM > 1 in at least one sample) in this species, and the expected number of recoded genes in each conservation group was calculated as the number of genes in this group multiplied by the expected probability of a gene being recoded. The significance level for the difference between observed and expected numbers in each conservation group was estimated by a two-tailed binomial test.

### Functional annotation and enrichment analysis of recoded genes

GO annotations for the protein-coding genes were downloaded from Ensembl (*Caenorhabditis elegans*, *Ciona savignyi*, *Danio rerio* and *Homo sapiens*) or Ensembl Metazoa (*Mnemiopsis leidyi*, *Amphimedon queenslandica*, *Drosophila melanogaster*, *Drosophila simulans*, *Crassostrea gigas*, *Octopus bimaculoides*, *Nematostella vectensis* and *Strongylocentrotus purpuratus*) via the BioMart function. For *Hydra vulgaris*, *Aplysia californica*, *Acromyrmex echinatior*, *Ptychodera flava* and *Branchiostoma belcheri* that do not have publicly available GO annotations, we first aligned all the proteins of these species to the UniProt database (release-2019_04) using BLASTP (blast-2.2.26) with parameters *-F F -e 1e-5*. Then the best hit of each query gene was retained based on its BLASTP bit score, and the GO annotations of this best hit was assigned to the query gene.

GO enrichment analysis was conducted for genes with at least one recoding site of which the mean editing level across samples > 0.1, or the editing event shared by at least two samples, in order to reduce the influence of nonadaptive recoding sites that are likely the by-products of promiscuous ADAR activity. Hypergeometric tests were employed to examine whether the recoded genes of a species was enriched in a specific GO term in relation to background genes as previously described ^31^, by comparing the number of recoded genes annotated to this GO term, the number of recoded genes not annotated to this GO term, the number of background genes (i.e. the protein-coding genes with RPKM > 1 in at least one sample after excluding the recoded genes in the species) annotated to this GO term, and the number of background genes not annotated to this GO term. *P*-values were adjusted for multiple testing by applying FDR ^69^, and the GO terms with adjusted *p*-values < 0.05 in at least three species (Note: GO terms shared by *D. melanogaster* and *D. simulans* were only counted once here) were considered as the general functional categories preferred by metazoan recoding editing.

### Identification of conserved recoding events shared by multiple species

To identify recoding events shared by two or more species, we first identified the orthologous groups of genes (i.e. gene families) from the seventeen metazoan species with reliable RNA editing using OrthoFinder (v2.2.7) ^84^ with default parameters. For the gene families that contained recoded genes from multiple species, we aligned the protein sequences of the recoded genes using MUSCLE (v3.8.31) ^85^ with parameter *-maxiters 1000* and filtered poorly aligned positions using Gblocks (v0.91b) ^86^. Next recoding events occurring in the same position in the alignments and causing the same amino acid changes among at least two species were identified as conserved recoding events. Recoding events only shared by *D. melanogaster* and *D. simulans* were removed. Only recoding sites in which the mean editing levels were no less than 0.1 across samples of a species, or were shared by at least two samples, were used in this analysis. The complete list of recoding events shared by multiple species was presented in Supplementary Table 5.

### Data and code availability

The raw sequencing reads generated in this study are deposited in NCBI under the BioProject accession PRJNA557895 and are also deposited in the CNGB Nucleotide Sequence Archive (CNSA) with accession number CNP0000504 (https://db.cngb.org/cnsa/). RNA-editing sites, refined gene annotations and repeat annotations used in this study are deposited in the figshare repository under the link https://doi.org/10.6084/m9.figshare.10050437. Codes are available upon request.

## Acknowledgements

We are grateful to Nicole King (University of California, Berkeley, USA) for providing the frozen stocks of *S. rosetta* and *M. brevicollis* and the protocols for starting and maintaining the cultures, Bernard Degnan and Kathrein E. Roper (University of Queensland, Australia) for providing the biopsies of *A. queenslandica*, Leo W. Buss (Yale University, USA) for providing the starter culture of *T. adhaerens* and the protocol for maintaining the culture, Robert E. Steele (University of California, Irvine, USA) for providing the *H. vulgaris* samples, Ulrich Technau (University of Vienna, Austria) for providing the *N. vectensis* samples, Xiaotong Wang (Ludong University, China) for providing the *C. gigas* samples, Qi Zhou (Zhejiang University, China) for providing the *D. melanogaster* and *D. simulans* samples, Bo Dong (Ocean University of China) for providing the *C. savignyi* samples, and Changwei Shao for providing the *D. rerio* samples. This work was supported by the National Natural Science Foundation of China (No. 31501057), the Science, Technology and Innovation Commission of Shenzhen Municipality (No. JCYJ20170817150239127 and JCYJ20170817150721687), the Strategic Priority Research Program of Chinese Academy of Sciences (No. XDB31020000 and XDB13000000), a Lundbeck Foundation Grant (R190-2014-2827) and a Carlsberg Foundation grant (No. CF16-0663) to G.Z., and a European Research Council Consolidator Grant (ERC-2012-Co -616960) to I.R.-T.

## Author Contributions

Q.L. and G.Z. conceived the study; M.T.P.G. and Q.L. coordinated the sample collection from different labs around the world; N.L. and L.G. conducted lab work for the culture and collection of *T. adhaerens* samples; M.D.M. conducted lab work for the culture and DNA/RNA extraction of *S. rosetta* and *M. brevicollis*; M.A.S. and I.R.-T. conducted lab work for the culture and DNA/RNA extraction of *S. arctica and C. owczarzaki*; M.Q.M. collected the *M. leidyi* samples and performed DNA/RNA extraction; Y.-H.S. and J.-K.Y. collected the *S. purpuratus*, *P. flava* and *B. belcheri* samples and performed dissection for *S. purpuratus*; X.Z. managed library construction and sequencing of all species; P.Z., J.L., H.Y., Y.Z., Q.G., H.T. and X.Z. performed bioinformatic analyses under the supervision of Q.L.; G.Z., N.L., I.R.-T., M.Q.M. and J.-K.Y. contributed reagents and materials; Q.L. and G.Z. wrote the manuscript with the inputs from all authors. All authors read and approved the final manuscript.

---

**Supplementary Figure 1.**
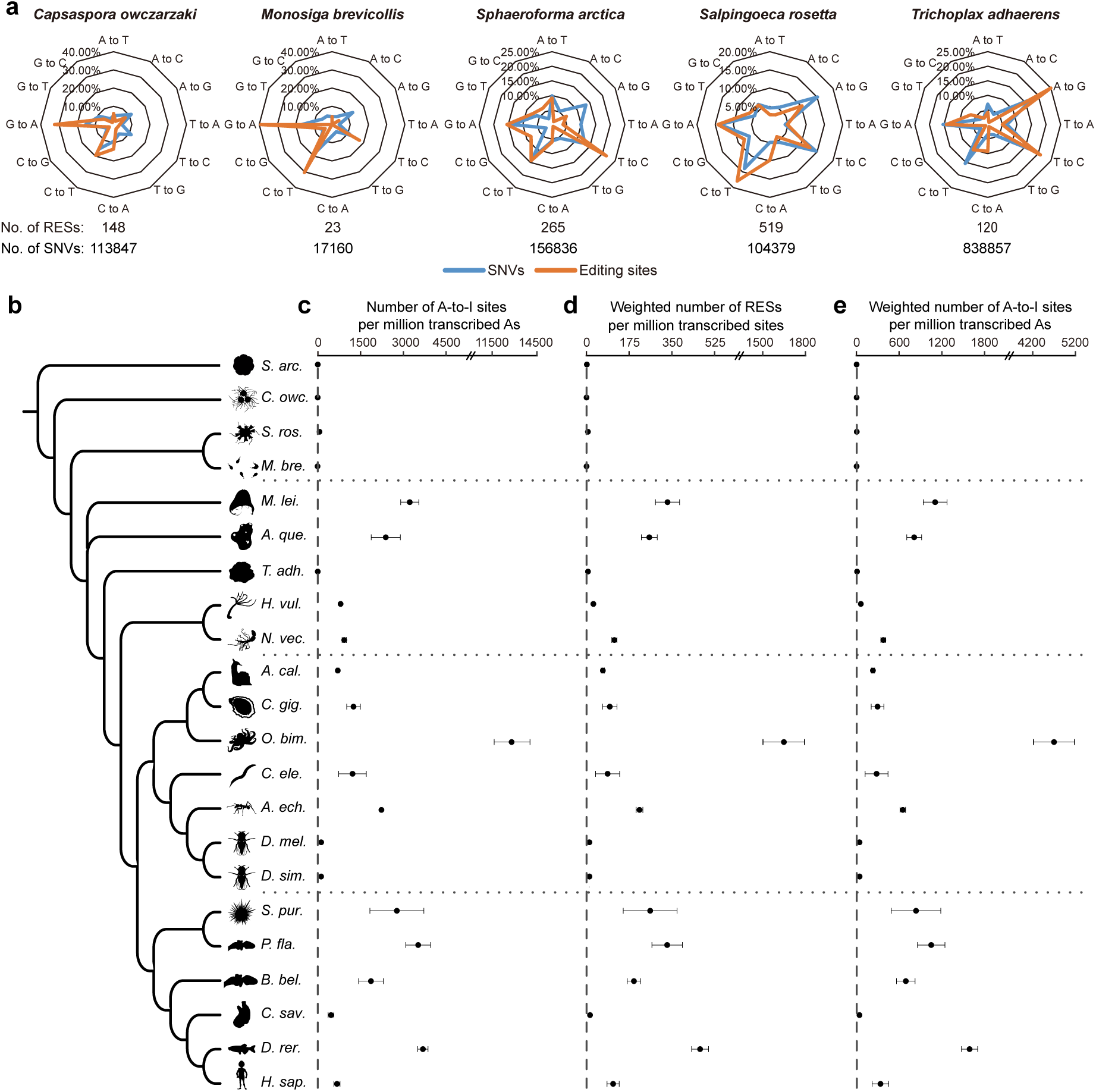
The origin and variation of RNA editing in metazoans. (**a**) The frequency of each type of nucleotide substitution among all potential RNA-editing sites (RESs) and genetic single nucleotide variants (SNVs) identified in the five species without *ADARs*. The types of nucleotide substitutions for RESs and SNVs were both inferred according to the genotypes present in the plus strand of the reference genome in this analysis. That is, an A-to-G RES from the minus strand of the reference genome was regarded as a T-to-C substitution, while substitution types of RESs from the plus strand remained unchanged. The RESs and SNVs from different samples of the same species were first combined according to their genomic locations, respectively, before the frequency calculation. (**b**) The phylogeny of the 22 species examined in this study. The topology of the phylogenetic tree is derived according to previous reports ^1, 2^. (**c**) The occurrence rate of A-to-I editing in each species, which was calculated as the number of A-to-I sites divided by the number of transcribed adenosine sites (RNA depth ≥ 2X) then multiplied by one million. (**d**) The editing-level-weighted occurrence rate of RNA editing in each species, which was calculated as the summed editing level for all RNA-editing sites with RNA depth ≥ 10X divided by the number of transcribed genomic sites with RNA depth ≥ 10X then multiplied by one million. (**e**) The editing-level-weighted occurrence rate of A-to-I editing in each species, which was calculated as the summed editing level for all A-to-I sites with RNA depth ≥ 10X divided by the number of transcribed adenosine sites with RNA depth ≥ 10X then multiplied by one million. Error bars in panels **c**, **d** and **e** represent s.d. across samples.

**Supplementary Figure 2.**
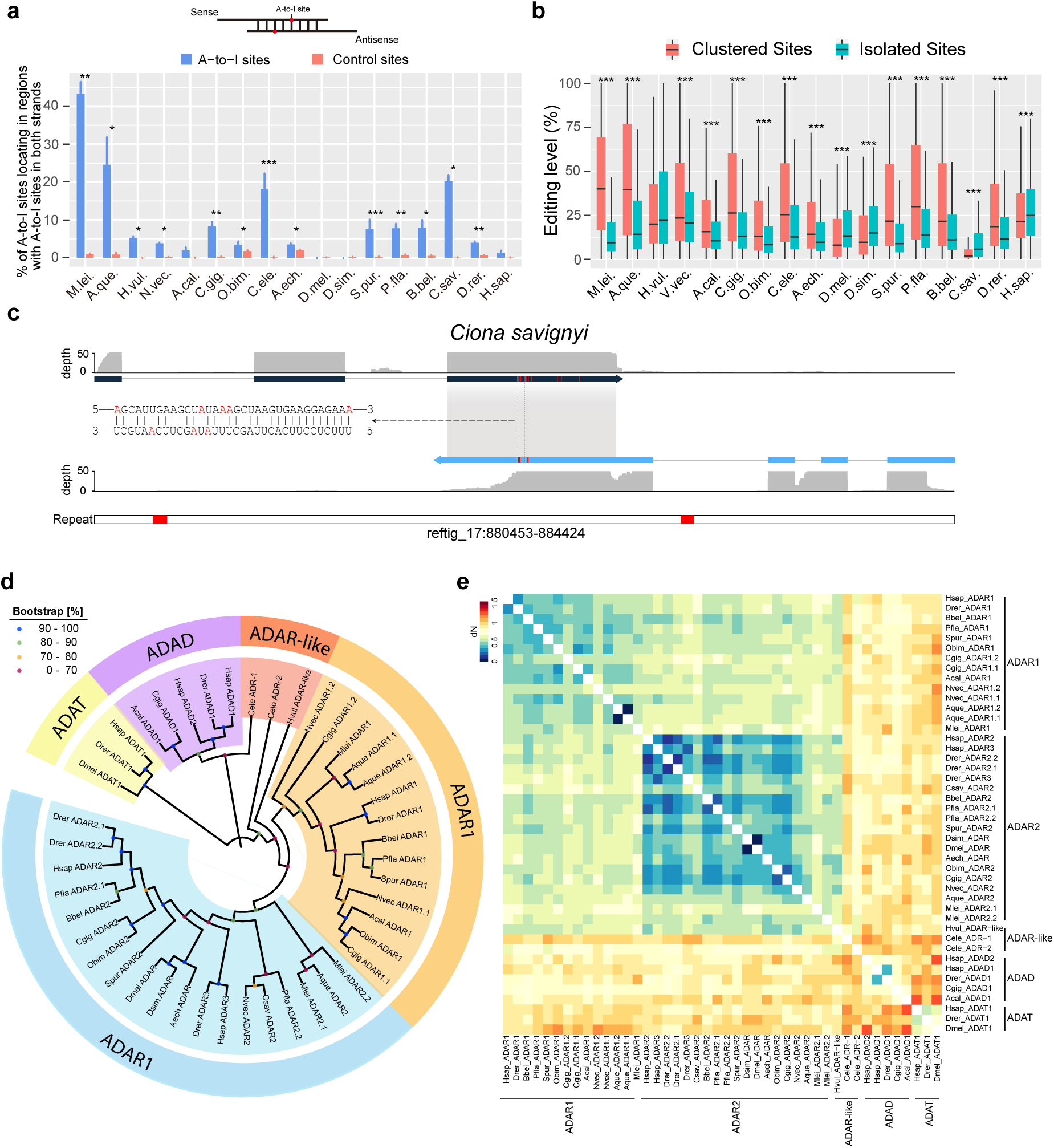
The structure and sequence preferences of A-to-I editing in metazoans. (**a**) The proportion of A-to-I editing sites locating in regions targeted by RNA editing on both strands, measured as the proportion of sites locating in a region (± 50 nt centered on the edited adenosine) with at least one A-to-I editing site found on the opposite strand. Control sites were randomly selected transcribed adenosines with the same number and comparable RNA depth of the A-to-I editing sites in each sample of each species (see Methods). The asterisks indicate significance levels estimated by two-tailed paired t-tests with “*” representing *p* < 0.05, “**” < 0.01 and “***” < 0.001. (**b**) The comparison of editing levels for the clustered and isolated A-to-I editing sites in each species. The asterisks indicate significance levels estimated by two-tailed Wilcoxon signed-rank tests with “*” representing *p* < 0.05, “**” < 0.01 and “***” < 0.001. (**c**) An example of a sense-antisense transcript pairing in *Ciona savignyi*, showing the RNA coverage of both transcript models, the pairing region (grey shadow), the A-to-I editing sites on both transcripts (red vertical bars), the detail view of a 34 nt regions (dashed box) with edited adenosines highlighted in red, and the distribution of repeats in this region (red boxes in the bottom track). (**d**) The phylogenetic trees of the deamination domains of ADARs using the neighbor-joining (NJ) method with MEGA7. Protein sequences of the deamination domains were aligned with ClustalW implemented in MEGA7. The deamination domains of ADAT1 from *D. melanogaster*, *D. rerio* and *H. sapiens* were selected as the outgroups. All protein sequences used for the phylogenetic analyses are present in Supplementary Table 3. (**e**) Pairwise nonsynonymous substitution rates (*dN*) for any pair of ADs in panel **d** using PAML (v4.9i) with the Yang & Nielsen (2000) method.

**Supplementary Figure 3.**
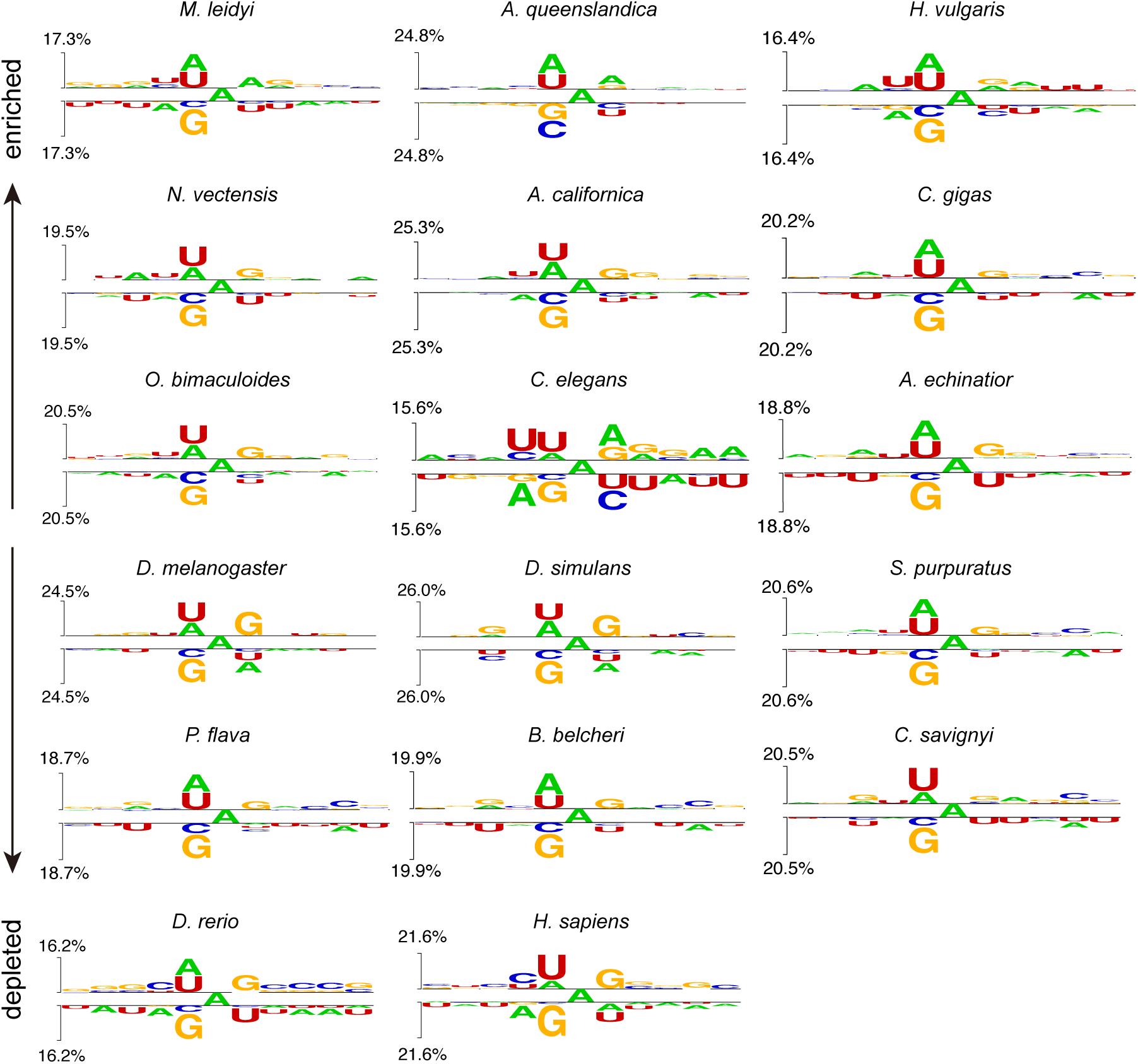
Neighboring nucleotide preference of edited adenosines. The neighboring nucleotide preference (± 5 nt) of the edited adenosines in each species was estimated in comparison to the unedited transcribed adenosines within ± 50 nt of the edited adenosines, by the Two Sample Logo software ^3^. Nucleotides were plotted using the size of the nucleotide that was proportional to the difference between the edited and unedited datasets, with the upper part presenting enriched nucleotides in the edited dataset and lower part presenting depleted nucleotides.

**Supplementary Figure 4.**
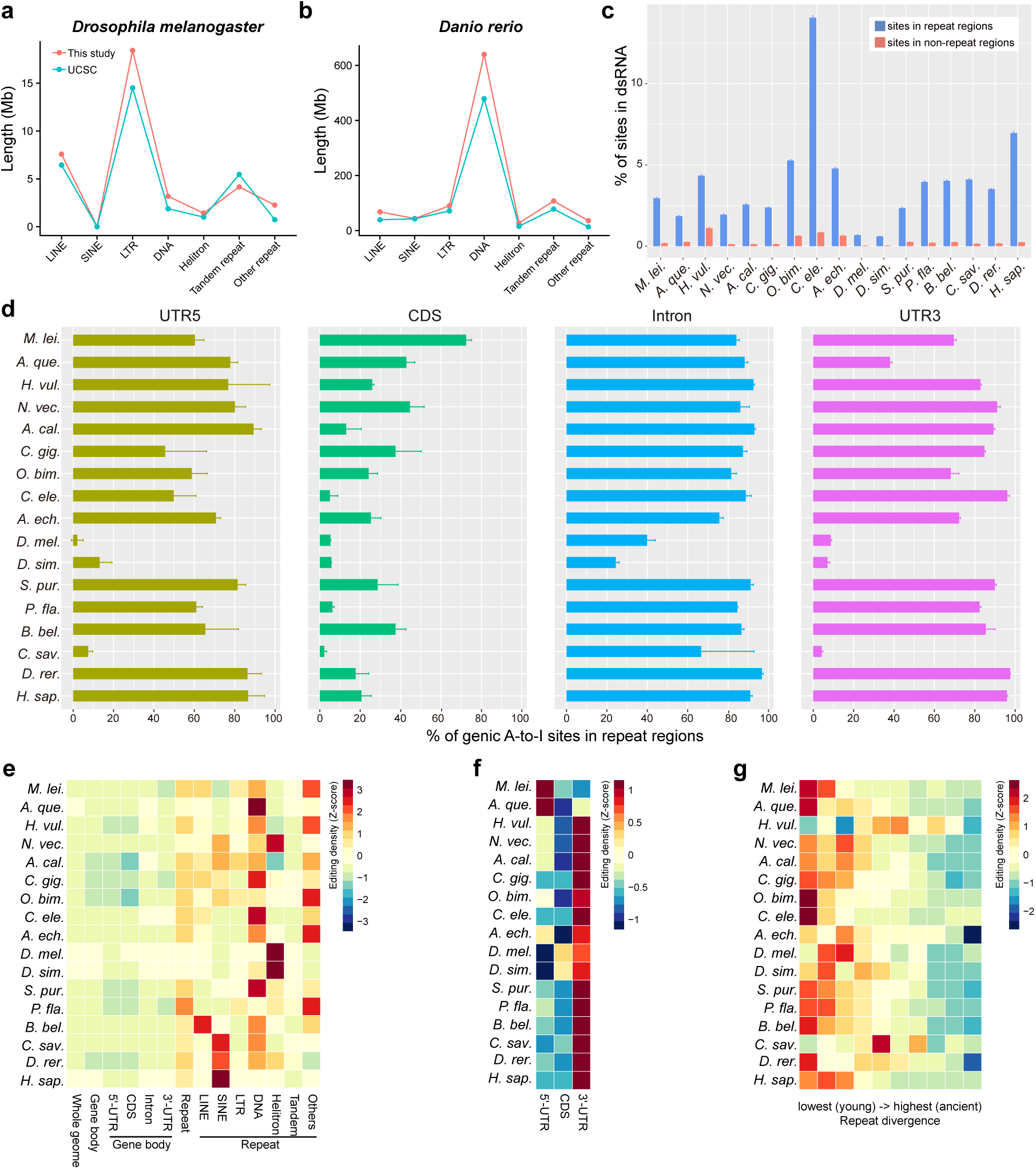
The primary genomic targets of metazoan A-to-I editing. (**a-b**) The non-redundant lengths of different repeat families in *D. melanogaster* (**a**) and *D. rerio* (**b**), according to the annotations generated in this study (see Methods) and those from UCSC, showing the good consistency between these two annotation results. (**c**) The potentials of repeat and non-repeat regions to form dsRNA in each species, measured as the ratios of repeat and non-repeat derived genomic sites locating in regions that could find a reverse-complement alignment in nearby regions (see Methods). *P*-values were estimated by Monte Carlo simulations (100 times) and < 0.01 for all species. (**d**) The percentage of genic A-to-I editing sites locating regions annotated as repeats. Genic editing sites were defined as editing sites locating in protein-coding gene related elements including 5’-UTR, CDS, intron and 3’-UTR. (**e**) Comparison of editing-level-weighted editing density across different genomic elements in each species. The weighted editing density of an element was calculated as the summed editing level of A-to-I editing sites (RNA depth ≥ 10X) locating in this element divided by the number of transcribed adenosines (RNA depth ≥ 10X) in this element. (**f**) Comparison of editing-level-weighted editing density across 5’-UTR, CDS and 3’-UTR. Note the different scales used between panel **e** and **f** in order to show the difference of editing density among genic elements. (**g**) The negative correlation between the sequence divergence and editing-level-weighted editing density of repetitive elements.

**Supplementary Figure 5.**
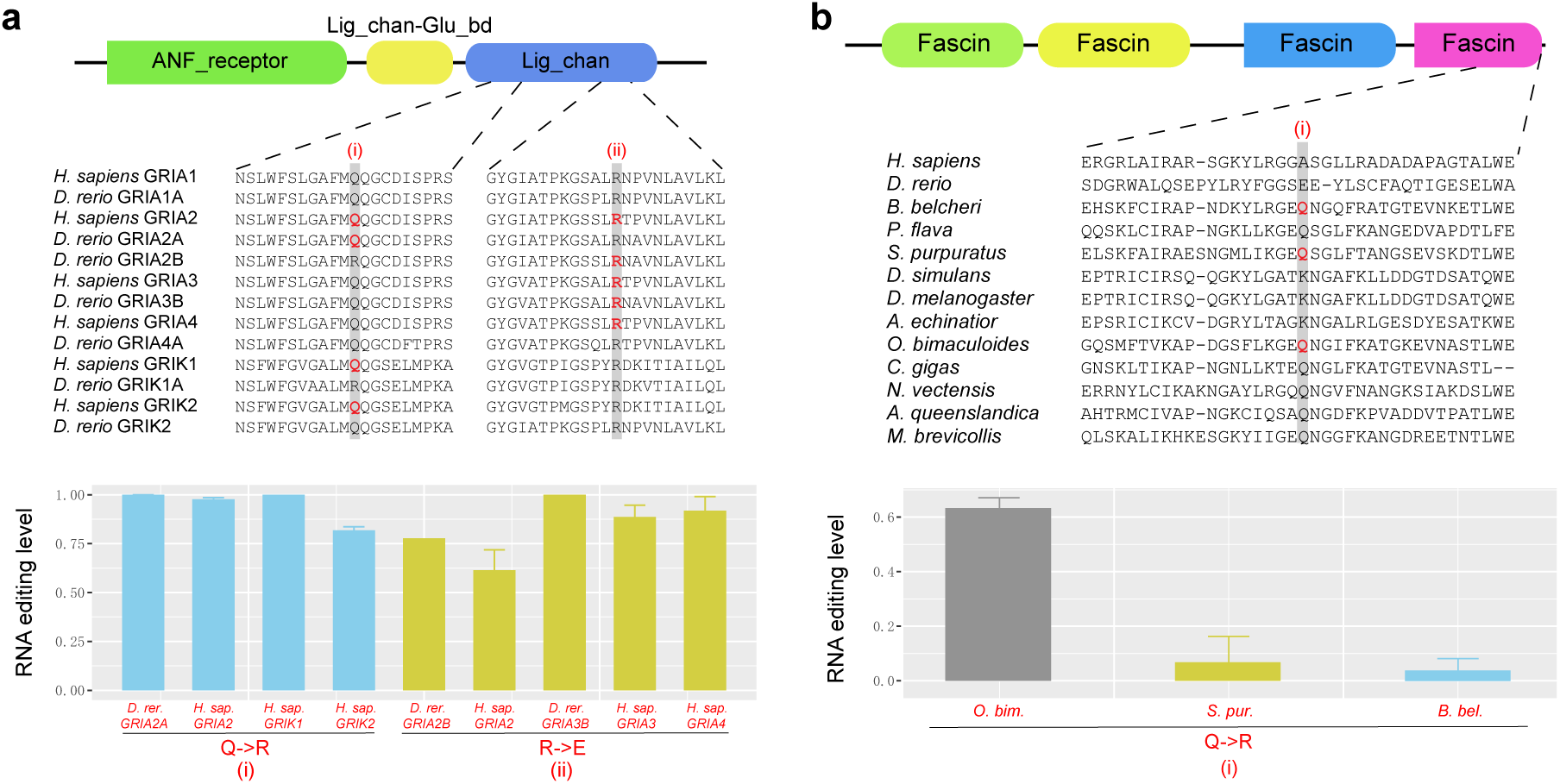
Functional preference of recoding editing in metazoans. (**a**) The convergent evolution of two recoding editing events in the glutamate ionotropic receptors in human and zebrafish. The upper part shows the domain organization of the human GRIA2 protein (NP_001077088.1) annotated by the Conserved Domain Database (CDD) of NCBI. The middle part shows the multiple sequence alignments of the two regions containing the shared recoding sites. Recoding sites were highlighted by red color and the positions of conserved recoding events were highlighted by gray shadow. The lower part shows the editing levels of the recoding events in each species. (**b**) The convergent evolution of a recoding editing event in fascin (an actin filament-bundling protein), which was shared by the octopus *O. bimaculoides*, the sea urchin *S. purpuratus* and the lancelet *B. belcheri*. The upper part shows the domain organization of lancelet fascin-like protein (XP_019642030.1) annotated by the CDD of NCBI. The middle and lower part are the same as panel **a**. Error bars in panels **a** and **b** represent s.d. across samples.

